# “Parvalbumin interneuron activity induces slow cerebrovascular fluctuations in awake mice”

**DOI:** 10.1101/2024.06.15.599179

**Authors:** Adiya Rakymzhan, Mitsuhiro Fukuda, Takashi Daniel Yoshida Kozai, Alberto Luis Vazquez

**Affiliations:** Department of Bioengineering, University of Pittsburgh, Pittsburgh, PA, United States of America; Center for the Neural Basis of Cognition, University of Pittsburgh and Carnegie Mellon University, Pittsburgh, PA, United States of America; Department of Radiology, University of Pittsburgh, Pittsburgh, PA, United States of America; Center for Neuroscience, University of Pittsburgh, Pittsburgh, PA, United States of America; McGowan Institute for Regenerative Medicine, University of Pittsburgh, Pittsburgh, PA, United States of America; NeuroTech Center, University of Pittsburgh Brain Institute, Pittsburgh, PA, United States of America

**Keywords:** blood flow, functional magnetic resonance imaging, optogenetics, parvalbumin, vascular fluctuations

## Abstract

Neuronal regulation of cerebrovasculature underlies brain imaging techniques reliant on cerebral blood flow (CBF) changes. However, interpreting these signals requires understanding their neural correlates. Parvalbumin (PV) interneurons are crucial in network activity, but their impact on CBF is not fully understood. Optogenetic studies show that stimulating cortical PV interneurons induces diverse CBF responses, including rapid increases, decreases, and slower delayed increases. To clarify this relationship, we measured hemodynamic and neural responses to optogenetic stimulation of PV interneurons expressing Channelrhodopsin-2 during evoked and ongoing resting-state activity in the somatosensory cortex of awake mice. Two-photon microscopy (2P) Ca2+ imaging showed robust activation of PV-positive (PV+) cells and inhibition of PV-negative (PV-) cells. Prolonged PV+ cell stimulation led to a delayed, slow CBF increase, resembling a secondary peak in the CBF response to whisker stimulation. 2P vessel diameter measurements revealed that PV+ cell stimulation induced rapid arterial vasodilation in superficial layers and delayed vasodilation in deeper layers. Ongoing activity recordings indicated that both PV+ and PV-cell populations modulate arterial fluctuations at rest, with PV+ cells having a greater impact. These findings show that PV interneurons generate a complex depth-dependent vascular response, dominated by slow vascular changes in deeper layers.

## Introduction

Disruptions in synchronized neural oscillations are linked to a wide range of cognitive impairment disorders such as schizophrenia or Alzheimer’s disease. These changes in neural network function occur early and serve as potential disease biomarkers [1–2]. In humans, these neural oscillations have been reported using electroencephalography and functional magnetic resonance imaging (fMRI) [3–7]. Known for its non-invasiveness and high sensitivity, fMRI is commonly used to monitor brain activity in humans, where the signal emerges from changes in cerebral blood flow (CBF) produced by neural activity [8]. However, interpreting these slow CBF signals is challenging without a more complete understanding of their neural correlates, as CBF is only a secondary measurement of neural activity.

Neurons actively modulate local blood supply by controlling the nearby vascular tone of arteries and arterioles enwrapped by smooth muscle cells that dilate and constrict the vessel [9]. However, the regulation of CBF by neurons remains elusive, as multiple cell types show different contributions to vascular changes. Previous studies in rodents show that excitatory neurons, which make up the majority of neurons in cortex, are largely activated during sensory stimulation and directly engage fast vascular responses [10–11]. Inhibitory interneurons, which comprise only 20% of all neurons, appear to change CBF independently of the activity of excitatory neurons [12–22]. Fast-spiking parvalbumin (PV) neurons are of particular interest since they are the most abundant of all inhibitory neurons in the cortex and their firing activity is considered to be the major source for generating gamma oscillations [12, 23–24]. Given the high hemodynamic correlation with neural gamma power in fMRI studies conducted in non-human primates and rodents and high vascular length correlation with PV neuron density, an important question to investigate has been whether blood flow is directly modulated by PV neurons [25–28].

Previous in vivo optogenetic studies reported conflicting effects of PV cell activation on CBF [13, 19–22]. Driving PV cells in brain slices constricted penetrating arterioles [15]. This observation agrees with the results from an in vivo wide-field optical imaging (OIS) study, where optogenetic stimulation of PV cells in awake PV-ChR2 mice induced an immediate CBF decrease, while in anesthetized animals, the same optogenetic stimulus induced a fast CBF increase [21]. A delayed slow CBF increase was observed in response to optogenetic stimulation in both anesthetized transgenic PV-ChR2 mice recorded with laser speckle contrast imaging (LSCI) and awake mice expressing ChR2 in PV cells through viral transfection recorded with Laser Doppler flowmetry (LDF) [19–20]. In an anesthetized mouse fMRI study, positive blood oxygen level-dependent (BOLD) signals were observed in the stimulation site upon optogenetic activation of PV cells, while the surrounding deeper area showed negative BOLD responses [13]. A recent optogenetic study in anesthetized mice, combining both fMRI and OIS, demonstrated a biphasic response at the stimulation site – fast vasoconstriction followed by slow vasodilation [22]. The discrepancies in the literature have been attributed to differences in the measured cortical depth (superficial vs. deep layers), stimulation protocols (short vs. long stimulation period), brain states (anesthetized vs. awake), or the method of expressing ChR2 (virus injection vs. transgenic mouse). A systematic assessment of the impact of PV neuron activity evoked by a range of stimulation parameters on vascular tone as well as on network activity in awake mice is essential to determine relevant contributions of these different vascular response features. Two-photon microscopy provides a suitable tool to examine these contributions at the cellular level as a function of depth with great specificity. More importantly, establishing the contributions of these relationships, or response features, during resting-state (stimulus-free) conditions has yet to be elucidated.

In this study, we aimed to (1) systematically assess the impact of PV neuron activity evoked by a range of stimulation parameters on CBF responses and vessel tone in awake mice, and (2) determine the contribution of ongoing PV neuron activity on spontaneous vascular fluctuations. We first hypothesized that driving PV neurons induces a slow increase in arterial diameter (vasodilation). To test this, we replicated our previous study, recording CBF using laser Doppler flowmetry (LDF) in awake mice expressing ChR2 in PV neurons in the barrel cortex [19]. We delivered light stimuli of various durations, frequencies, and pulse widths. Subsequently, employing the same stimulation paradigm, we measured neuronal (PV+ and PV-) calcium activity and nearby vasculature across cortical depth using two-photon microscopy (2P). We then conducted resting-state 2P recordings to determine whether fluctuations in arteriolar diameter can be preferentially predicted by PV neuron activity.

## Materials and methods

### Animal surgery for awake head-fixed experiments

We used 13 adult transgenic mice (male n=10, female n=3; Jackson Laboratories, Bar Harbor, ME USA) expressing recombinase in PV neurons (PV; strain=B6.129P2-Pvalb(tm1-cre)Arbr/J). At 8-10 weeks of age, we used viral transduction to express channelrhodopsin-2 (ChR2) by injecting cre-dependent AAV encoding ChR2-eYFP (AAV-EF1a-DIO-ChR2-eYFP; University of North Carolina Vector Core, Chapel Hill, NC USA) in the primary somatosensory cortex (S1; Figure 1A-B). To drive the expression of calcium (Ca^2+^) reporter in neurons under the Synapsin promoter, mice were injected with AAV-Syn-RCaMP1a (Addgene, Watertown, MA USA) in the same region of S1 (Figure 1A, C). For the control group, in addition to experiments in non-expressing cortical regions in a subset of PV-cre mice expressing ChR2, we used 2 male adult PV-cre mice injected with AAV-EF1a-DIO-tdTomato (University of North Carolina Vector Core, Chapel Hill, NC USA) and AAV-Syn-GCaMP7f (Addgene, Watertown, MA USA) and 2 male adult mice expressing Ca^2+^ indicator jRGECO1a in cells under the Thy1 promoter (Thy1-jRGECO1a; strain: Thy1-jRGECO1a-WPRE, line GP8.31 with stock number 030527). Each animal underwent a surgery to implant a head holder and cranial window for optical access (Figure 1A). Mice were anesthetized with ketamine (75 mg/kg) and xylazine (10 mg/kg). A 4-mm craniotomy was performed over S1 and re-sealed with a glass coverslip, and a custom head-plate was fixed to the skull. For detailed surgical procedures, refer to [19]. Mice were given at least 4 weeks to recover post-surgery and before imaging. During the recovery period, mice were acclimated to head fixation on a treadmill (1 hour in their cage followed by 1 hour under the microscope for 5 days). Imaging occurred when the mice were 3-8 months old. All procedures followed an approved protocol by the University of Pittsburgh Institutional Animal Care and Use Committee (IACUC) by the standards for humane animal care and use as set by the Animal Welfare Act (AWA), the National Institute of Health Guide for the Care and Use of Laboratory Animals and ARRIVE (Animal Research: Reporting in Vivo Experiments).

**Figure 1.**
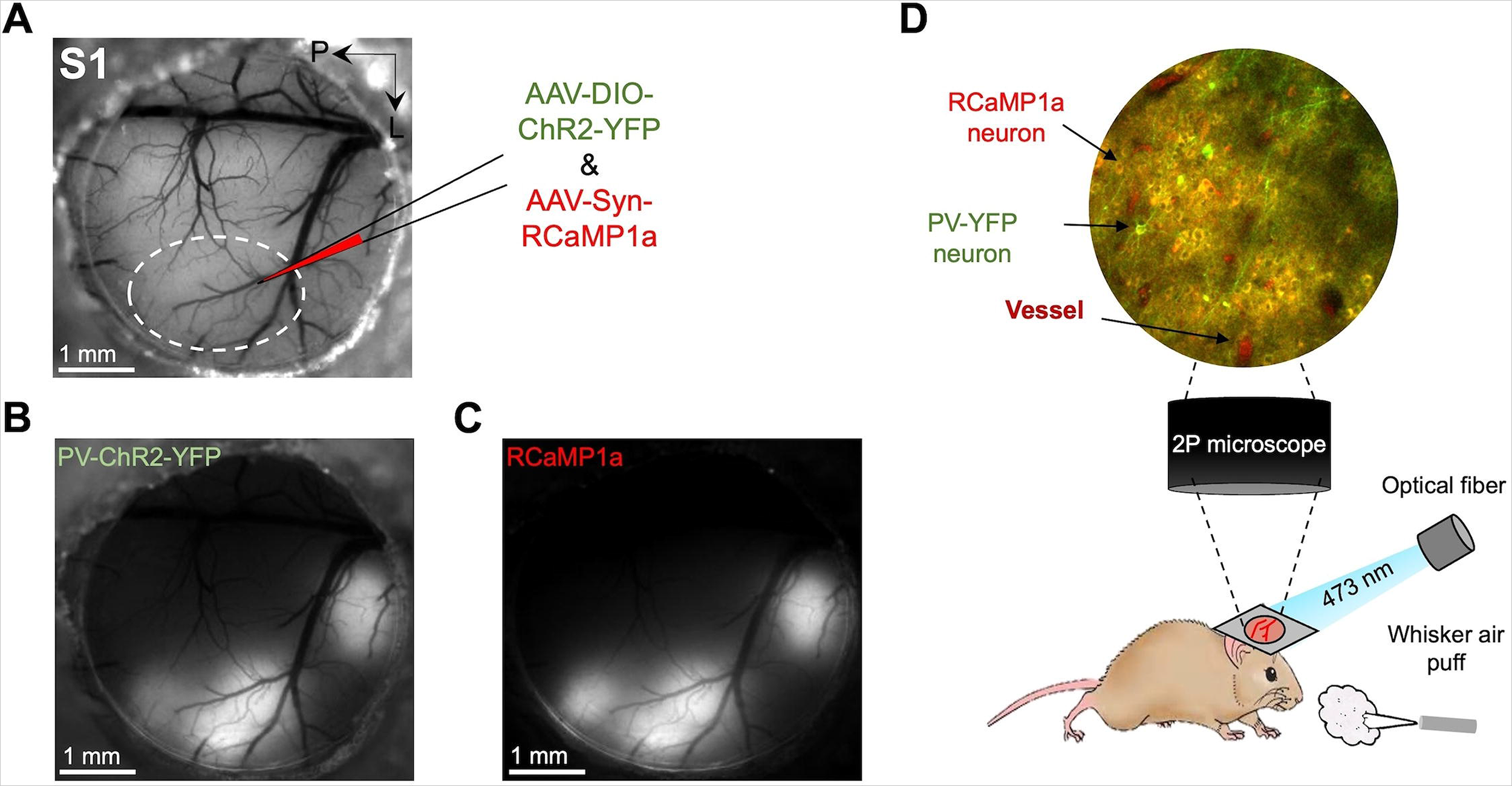
Schematic of the experimental setup. **A**. Cranial window implanted in PV-cre mouse S1 where we injected optogenetic protein (AAV-DIO-ChR2-YFP) and calcium reporter (AAV-Syn-RCaMP1a) viruses. Surface fluorescence microscope image showing the regions expressing: **B**. ChR2-YFP and **C**. RCaMP1a. **D**. Mouse was placed under a 2P microscope where neurons expressing RCaMP1a, including PV-YFP cells, and vessels were visualized. The optical fiber probe and air puff were placed for optogenetic stimulation and whisker stimulation experiments, respectively.

### Stimulation

Optogenetic stimulation was achieved using a 473 nm power-adjustable (max. 50 mW) TTL-controlled laser diode unit (CrystaLaser Inc., Reno, NV USA) connected to an optical fiber (S-405XP, ThorLabs Inc., Newton, NJ USA) with a core diameter of 125 μm placed at a 30° on the cover glass (Figure 1D, Figure 2A). With such a fiber placement, the intensity of the light stimulus was calibrated to 1 mW using an optical power meter, sufficient to induce neuronal activation across cortical depth [11]. Light stimulus pulse duration (2, 10, or 30 ms) and frequency (5, 20, or 40 Hz) were varied to modulate evoked activity and examine its impact on hemodynamic responses. For whisker stimulation, air puff pulses (50 ms/pulse) were delivered at 5 Hz using a pressure injector set to 30 psi (Figure 1D, Figure 2A). Stimuli were delivered over a 1-second period every 30 s or 4 s every 40 s. At least 10 stimulation trials were collected for each parameter set.

**Figure 2.**
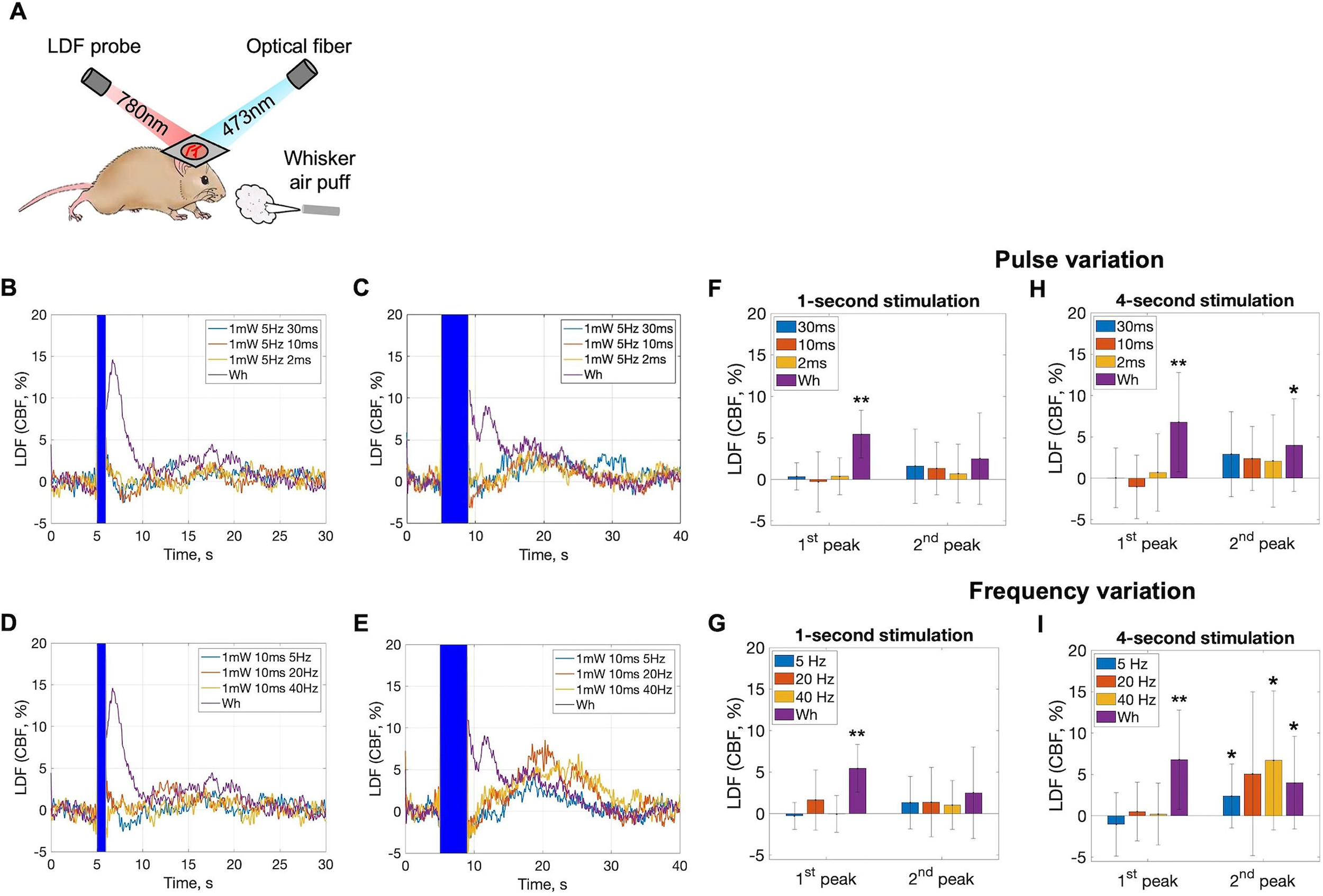
Optogenetic stimulation of PV neurons induces a slow increase in CBF measured with LDF. **A.** LDF probe for CBF measurements and an optical fiber for blue light delivery were placed on top of a glass coverslip in awake mice. Air puffs were delivered for whisker stimulation experiments. **B-C**. Time profiles of average CBF changes measured by LDF in response to optogenetic stimulation of PV-ChR2 cells at various pulse widths at 5 Hz for 1 s and 4 s, and **D-E**. at various frequencies with 10-ms pulse width, as well as whisker stimulation. The blue rectangle denotes the blue light stimulation period. **F**. Summary of average CBF changes in response to 1-second optogenetic stimulation at different pulse widths and **G**. at different frequencies. **H**. Summary of average CBF changes in response to 4-second optogenetic stimulation at different pulse widths and **I**. at different frequencies. Error bars denote standard deviation. Significant differences from baseline are denoted by * for p<0.05 or by ** for p<0.01. n=10-13 animals.

### Laser Doppler flowmetry

A fluorescence microscope (MVX-10, Olympus Inc., Japan) identified ChR2-YFP and RCaMP1a expression regions in S1 (Figure 1B). A Laser Doppler Flowmeter (LDF; Periflux 5000/411, Perimed AB, Jarfalla, Sweden) acquired CBF data sampled at 1000 Hz. The LDF probe, with a tip diameter of 450 μm and operating wavelength of 780 nm, was placed at 30° on the cover glass facing the expressing region, avoiding large surface vessels (Figure 2A). For control experiments, the optical fiber and LDF probe were positioned in a non-expressing cortical region.

### 2P imaging

Each mouse underwent at least 3 awake imaging sessions during the daytime with a week in between sessions. Before imaging, mice were intraperitoneally injected with Texas Red Dextran (3kDa, 5% w/w solution; Thermo Fisher Scientific, Waltham, MA USA) for blood flow contrast. Imaging was performed with an InSight X3 widely tunable ultrafast laser (Spectra-Physics, Milpitas, CA USA) on a two-photon fluorescence microscope (Ultima IV, Bruker Nano Inc., Billerica, MA USA) with a 16X water immersion objective lens, 0.80 NA (Nikon Inc., Tokyo, Japan) with a maximum field-of-view (FOV) of 800 × 800 μm (Figiure 1E). Two-photon excitation was performed at 1100 nm for imaging RCaMP1a and Texas Red Dextran up to 400 μm depth (layers 1-3) in S1. Imaging parameters were raster scanning rate at 2.4 ms/pixel; 332 × 332 μm FOV with 1.66 μm/pixel; 200 × 200 matrix; 5 Hz frame rate. The selection of the imaged region was based on four parameters, where the FOV includes: penetrating large vessels, RCaMP1a cells, PV-YFP cells, and barrel cortex.

### Data analysis

All data were analyzed using MATLAB (Mathworks, Natick, MA USA).

### CBF

Time series of CBF changes evoked by whisker or optogenetic stimulation were obtained from LDF data. The CBF time series were low-pass filtered with a 5-Hz rectangular cut-off and down-sampled to 10 Hz.

### Cell region of interest (ROI)

Active cells expressing Ca^2+^ reporter RCaMP1 were segmented from the average intensity 2P image using a custom-written function in MATLAB. The function segments cells based on correlation, identifying connected pixels based on temporal correlation with neighboring pixels. PV-positive (PV+) and PV-negative (PV-) neurons were identified from a fluorescence difference map produced by subtracting the average pre-stimulation baseline (average over 5 s prior stimulation period) from the average post-stimulation frame (average over 2 s following stimulation period). A cell was considered PV+ if its Ca^2+^ fluorescence change reached at least 5% during 2 s immediately following the stimulation at least 50% of the time. A cell was considered PV-if its Ca^2+^ fluorescence change dropped by at least 5% during 2 s immediately following the stimulation at least 50% of the time. The optogenetic stimulus covers portions of the images during stimulation, so we only considered pre- and post-stimulation images to extract ROIs. The final ROIs were qualitatively examined to match the morphological features of neurons. The Ca^2+^ fluorescence signal of each cell body was measured by averaging all pixels within the ROI. Ca^2+^ change traces were corrected for neuropil contamination by regressing out the first two principal components of the neuropil Ca^2+^ change trace.

### Diameter measurement

Vessel diameter was measured from 2P images as the full-width-at-half-maximum of the vessel cross-sectional intensity profile. Diameter changes in both stimulation and ongoing activity experiments were filtered using a Fermi filter with cutoff frequency of 1 Hz and transition band of 0.2 Hz. Slow vascular activity was isolated by filtering the vessel diameter change using a cutoff frequency of 0.06 Hz and transition band of 0.01 Hz.

### Time series

The CBF time series, neuronal Ca^2+^ traces, and diameter changes with common stimulation parameters were averaged and then normalized by their pre-stimulation baseline level (average over the 5 s preceding stimulation onset). The Ca^2+^ traces of individual neurons and the diameter changes of an individual vessel during the ongoing activity were detrended using a 5^th^-order polynomial and then normalized by their modes.

### Hemodynamic response function (HRF) fit

The HRF was defined by a gamma function with four parameters. The gamma fit parameters were determined using ‘lsqnonlin’ optimization routine in MATLAB to minimize the root mean squared error between the measured CBF response or measured vascular activity and neural input convolved with the gamma function. The estimated vascular activity was obtained for each neuron by convolving the Ca^2+^ fluorescence signal with HRF.

### CBF response estimation

To estimate the CBF response to whisker stimulation, the following model was employed, combining multiple HRFs:

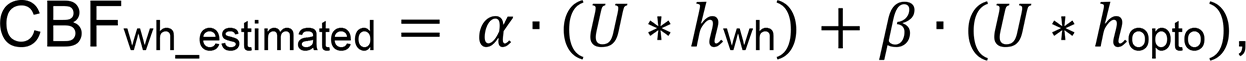

where *U* represents neural input, *h*wh represents the average HRF from whisker stimulation trials, *h*opto represents the average HRF from optogenetic stimulation trials, and α and β are coefficients calculated by solving a linear equation:

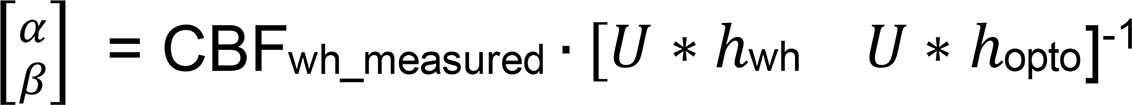

*Peak calculation:* We measured changes in CBF and vessel diameter over two locations: immediately after the stimulation offset and 4-22 s after stimulation offset, to capture fast and slow vascular response magnitudes. These changes were recorded from the time series. Optogenetic stimulus artifacts in LDF readings and 2P images prevented signal change calculations during stimulation, so changes were determined immediately after stimulation ended. The 1^st^ peak magnitude was calculated as the average response over 4 s starting from the peak time ([0 s; 4 s], where 0 s is the 1^st^ peak time). The 2^nd^ peak magnitude was calculated as the average response over 4 s around the peak time ([-2 s; 2 s], where 0 s is the 2^nd^ peak time). The 1^st^ peak time was determined as the time point with the highest amplitude of the absolute signal within 4 s immediately following the stimulation offset, considering both positive and negative peaks. The time of the 2^nd^ peak was determined as the time point with the highest amplitude of the signal within 18 s starting from the 4^th^ s following the stimulation offset, considering both positive and negative peaks. The change in Ca^2+^ fluorescence was measured as the average response over 2 s immediately following the stimulation offset.

### Statistical analysis

We assessed significant differences in response amplitude relative to the baseline or between sexes using a t-test (p<0.05 or p<0.01). To test relationships and address multiple comparisons, a linear mixed-effect model was employed for response amplitude as a function of pulse duration or frequency (p<0.05). A linear regression model was applied to the peak time of vessel diameter change as a function of cortical depth, considering significant relationships (p<0.05). For analyzing the correlation between measured and estimated vascular activity, Pearson’s linear correlation coefficient was computed using ‘corr’ function in MATLAB. The significance of correlation coefficients was tested using a t-test (p<0.05). We report average values ± standard deviation throughout the manuscript and figures.

## Results

### Optogenetic stimulation of PV neurons induces a slow increase in CBF

First, we replicated our previous experiments examining CBF responses to optogenetic stimulation in awake head-fixed PV-cre mice expressing ChR2 in PV neurons in the barrel cortex [19]. Using LDF, we recorded CBF while delivering photo stimuli of various durations, frequencies, and pulse widths (Figure 2A). On average, 1-second optogenetic stimulation of PV neurons did not elicit discernible CBF changes (Figure 2, B, D, and F-G). However, a 4-second optogenetic stimulation induced prominent 2^nd^ peaks, especially at higher frequencies. At 5 Hz, 20 Hz, and 40 Hz, the 2^nd^ peaks were 2.4 ± 3.9% at 18.3 s, 5.1 ± 9.9% at 19.2 s, and 6.7 ± 8.4% at 20.3 s after stimulus onset, respectively (Figure 2E). Statistical significance was observed for the 5-Hz (n=13; p<0.05; Figure 2I) and 40-Hz stimulations (n=10; p<0.05; Figure 2I). Although the average 2^nd^ peak in the 4-second stimulation increased with frequency, linear mixed-model analysis demonstrated no significant relationship between peak amplitude and stimulation frequency (p>0.39). Varying pulse widths did not significantly impact the CBF response (Figure 2B-C, F, H). The 1^st^ peak in the 5-Hz 10-ms 4-second stimulation appeared negative (Figure 2C, E), with an average amplitude of -1.0% ± 3.8% at 4.9 s after stimulus onset, but it was not significant (n=13, p=0.34; Figure 2H-I). Since experiments were conducted in animals of both sexes, we tested for differences in CBF responses between sexes and did not observe significant differences (6.9±3.6% vs. 6.1±6.7%, 10 males vs. 3 females, p=0.94), although our study was not designed or appropriately powered for this purpose. Therefore, from here on data were pooled across sexes. As previously shown, optogenetic stimulation did not cause CBF changes in ChR2 naïve animals (i.e. data collected from areas without ChR2 expression in experimental mice did not reveal any significant changes in CBF after optogenetic stimulation (Supplemental Figure 1) [19]. Consistent with our prior findings, these results confirm that prolonged (4 s) optogenetic stimulation of PV neurons induces a delayed, slow increase in CBF.

Both 1-second and 4-second whisker stimulations elicited typical fast CBF responses. The 1-second stimulation induced a 1^st^ peak of 5.4 ± 2.9% and a 2^nd^ peak of 2.5 ± 5.5% at 1.8 s and 14.4 s after stimulus onset, respectively (Figure 2B, D). For the 4-second stimulation, the 1^st^ peak was 6.8 ± 6.0%, and the 2^nd^ peak was 4.0 ± 5.6% at 5.3 s and 17.3 s after stimulus onset, respectively (Figure 2C, E). Statistical significance was observed for the 1^st^ peak in both durations (n=12; p<0.01; Figure 2F-I) and for the 2^nd^ peak only in the 4-second stimulation (n=12; p<0.05; Figure 2H-I). A peak observed behind the stimulation bar in the 4-second whisker stimulation experiment (Supplemental Figure 2) aligned with the 1^st^ peak in the 1-second whisker stimulation and was disregarded for consistency with optogenetic experiments. Overall, these experiments demonstrated that neurovascular coupling was maintained in these mice.

### Linear convolution model of the CBF response to whisker stimulation

Next, we estimated the CBF response to whisker stimulation using a linear convolution model to examine whether the sustained whisker response could be captured by prolonged PV neuron responses. The neural input was represented by a rectangular function corresponding to the stimulation period (Figure 3A-B) and a gamma function was used to estimate its corresponding response function (HRF). We averaged HRFs across all trials for 4-second optogenetic stimulation of PV cells (Figure 3C) and convolved this average HRF with the neural input. This yielded the estimated CBF response to 4-second optogenetic stimulation of PV cells with r^2^=0.74 between the measured and estimated CBF responses (Figure 3E). We repeated this process for 1-second whisker stimulation and obtained r^2^=0.81 (Figure 3D-F). Note that while the obtained model successfully captured the first peak, it failed to capture the second, smaller peak around 17 s. However, the model that combined both optogenetic and whisker stimulation HRFs (Figure 3G, I) captured both the first and second peaks in the CBF response to whisker stimulation (Figure 3H, J). This model estimated CBF responses with r^2^=0.92 and r^2^=0.69 to 1-second and 4-second whisker stimulations, respectively, and that is captured well the sustained (or delayed) response portion of the CBF response. This suggests that the sustained response to whisker stimulation might be due to PV neuron activity.

**Figure 3.**
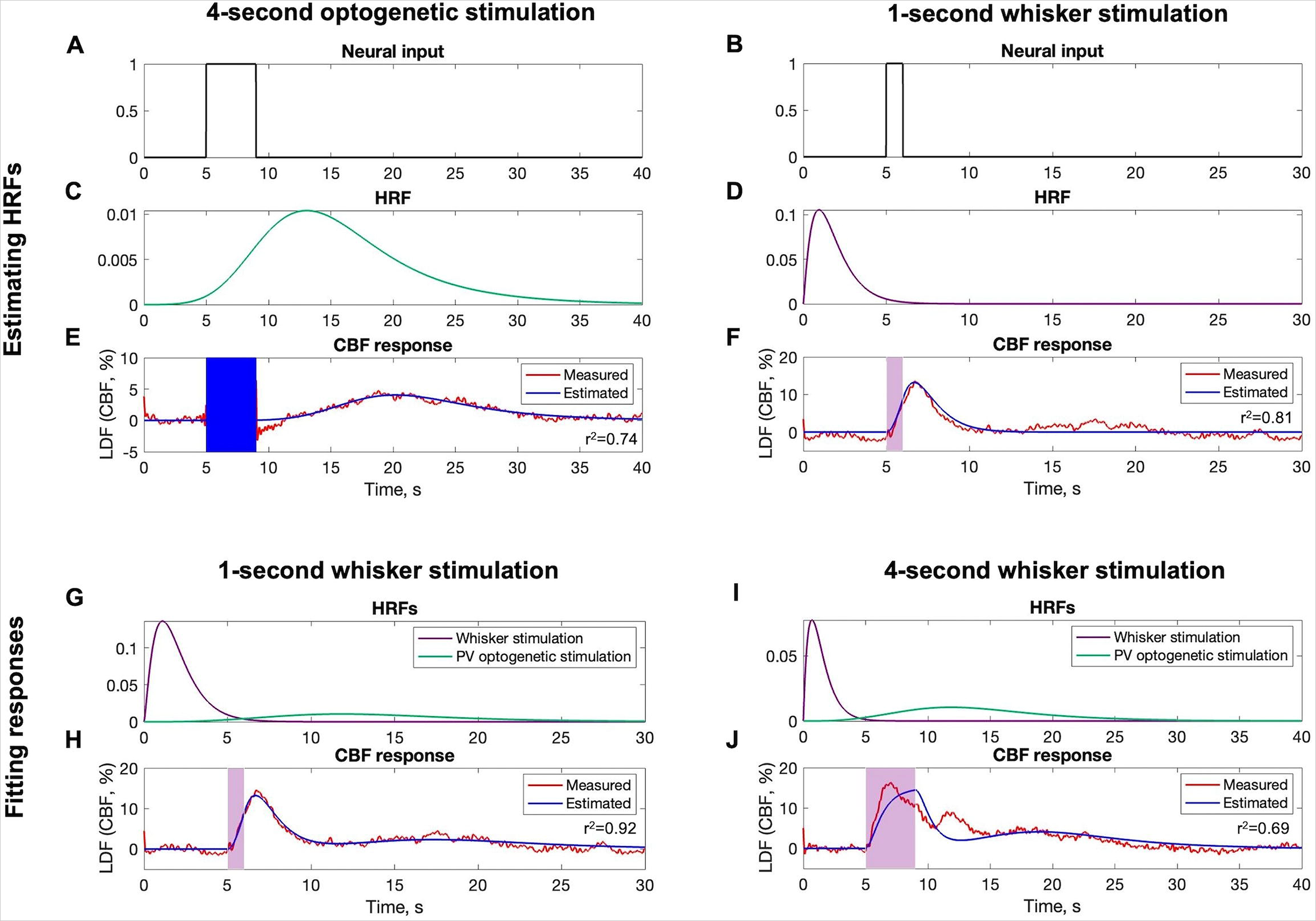
Estimation of the LDF-measured CBF response to whisker stimulation using linear convolution model. **A.** Neural input represented by a rectangular function for 4-second and **B**. 1-second stimulations. **C**. Average HRF obtained from 4-second optogenetic and **D**. 1-second whisker stimulation trials. **E**. The measured and estimated CBF responses to optogenetic and **F**. whisker stimulations. **G, I**. Average HRFs used to estimate **H.** CBF responses to 1-second and **J**. 4-second whisker stimulation. The purple semi-transparent rectangle denotes whisker stimulation period, the blue rectangle denotes optogenetic stimulation period. n=10-13 animals.

### Optogenetic stimulation activates PV+ neurons and inhibits PV-neurons

Next, we investigated the impact of optogenetic stimulation on PV neuron and network activity in awake head-fixed mice by 2P. Neuronal activity was recorded using the Ca^2+^ sensor RCaMP1a, while PV neurons expressing ChR2 were activated with blue light (473 nm). Two-photon images of intra-cortical sections about 400 μm below the cortical surface (deep layer 3) showed successful ChR2 expression in PV neurons, indicated by YFP fluorescence along cell membranes (Figure 4A) and RCaMP1a fluorescence in all neurons (Figure 4B) about one-month post-virus injection. Population neuronal activity was recorded with RCaMP1a in both PV+ and PV-cells during optogenetic stimulation of varying durations. Since pulse width variations did not affect the CBF response, we focused on varying only the stimulation duration (1 s and 4 s) and frequency (5 Hz, 20, Hz, 40 Hz). The Ca^2+^ fluorescence difference map immediately after stimulation revealed that 1-second optogenetic stimulation (5-Hz trains of 10-ms light pulse) activated some cells (PV+) and inhibited others (PV-) (Figure 4D). The Ca^2+^ fluorescence traces were extracted from PV+ and PV-cell masks (Figure 4E-F).

**Figure 4.**
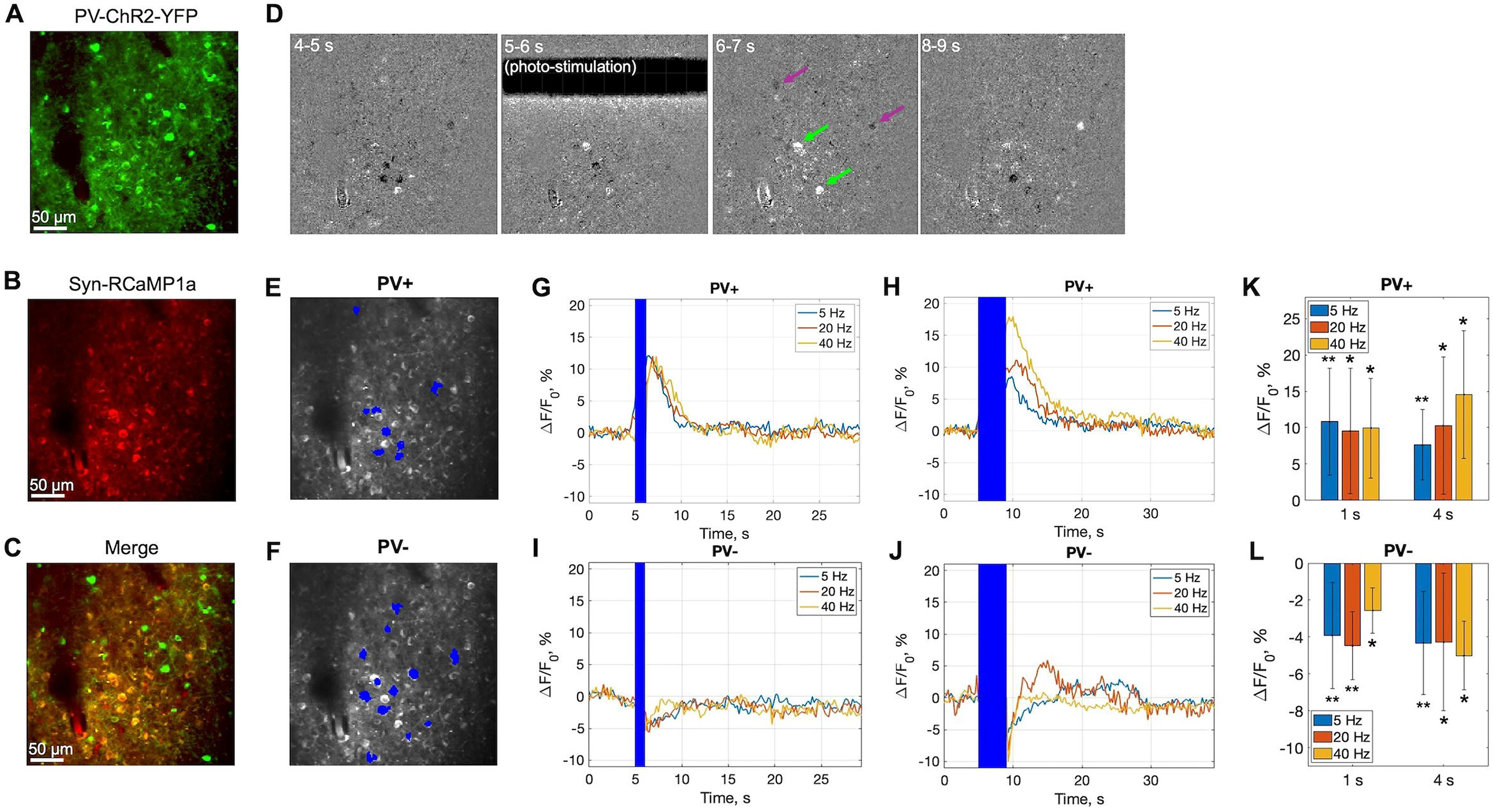
Optogenetic stimulation activates PV+ neurons and inhibits PV-neurons. **A.** 2P microscopy images of S1 intra-cortical sections showing the regions expressing ChR2-YFP, **B**. RCaMP1a, and **C**. color merge. **D**. An example of RCaMP1a difference maps at pre-stimulation (4-5 s), during stimulation (5-6 s), immediately after stimulation (6-7 s), and post-stimulation (8-9 s). In this particular example, a 10-ms optogenetic stimulus was delivered for 1 s at 5 Hz. Optogenetic stimulus, some activated and some inhibited cells are indicated by black bar, green and purple arrows, respectively. **E**. PV+ and **F**. PV-cell masks generated from various frequency optogenetic stimulation experiments. **G**. Time courses of RCaMP1a changes in PV+ in 1-s and **H**. 4-s optogenetic stimulation experiment. **I**. Time courses of RCaMP1a changes in PV-in 1-s and **J.** 4-s optogenetic stimulation experiment. **K**. Summary of average RCaMP1a changes at different frequencies in PV+ (n=4-10 animals) and **L**. in PV-cells (n=4-9 animals). Error bars denote standard deviation. Significant differences from baseline are denoted by * for p<0.05 or by ** for p<0.01.

On average, both 1-second and 4-second optogenetic stimulation robustly increased Ca^2+^ fluorescence in PV+ cells, indicating activation with light (Figure 4G-J). For 1-second stimulation, the average Ca^2+^ fluorescence response in PV+ cells was consistent across frequencies, with amplitudes of 10.8 ± 7.4%, 9.5 ± 9.66%, and 9.9 ± 6.9% (n=10, p<0.01; n=9, p<0.05; n=5, p<0.05) at 2.2 s, 2.2 s, and 2.4 s after stimulus onset for 5 Hz, 20 Hz, and 40 Hz, respectively (Figure 4K). In contrast, during 4-second stimulation, the average Ca^2+^ peak in PV+ cells appeared to increase with frequency. At 5 Hz, 20 Hz, and 40 Hz, the average Ca^2+^ fluorescence increase in PV+ cells was 7.6 ± 4.9%, 10.3 ± 9.5%, and 14.6 ± 8.8% (n=10, p<0.01; n=8, p<0.05; n=4, p<0.05) at 5.1 s, 5.5 s, and 5.7 s after stimulus onset, respectively (Figure 4K). Linear mixed-model analysis demonstrated no significant relationship between peak amplitude and stimulation frequency (p>0.31).

In PV-cells, the Ca^2+^ fluorescence drop in 1-second stimulation was consistent across frequencies with amplitudes of -3.9 ± 2.9%, -4.5 ± 1.8%, and -2.6 ± 1.23% (n=9, p<0.01; n=7, p<0.05; n=4, p<0.05) at 2.1 s, 2.1 s, and 1.95 s after stimulus onset for 5 Hz, 20 Hz, and 40 Hz, respectively (Figure 4L). The Ca^2+^ fluorescence drop in 4-second stimulation was consistent across frequencies but with larger amplitudes: -4.3 ± 2.8%, -4.3 ± 3.7%, and -5.0 ± 1.9% (n=8, p<0.01; n=7, p<0.05; n=4, p<0.05) at 5.0 s, 4.8 s, and 4.7 s after stimulus onset for 5 Hz, 20 Hz, and 40 Hz, respectively (Figure 4L). These results confirmed that optogenetic stimulation activates inhibitory PV+ neurons expressing ChR2, subsequently inhibiting some neurons labelled PV-.

### Optogenetic stimulation of PV neurons induces depth-dependent vascular responses

Previous studies have used techniques biased towards either superficial layers, such as IOS or LSCI, or with no depth information, such as LDF. To address this, we systematically assessed the vascular responses to optogenetic stimulation of PV neurons in both superficial and deeper cortical layers separately [19–22]. We concurrently imaged pial and penetrating vessels while optogenetically activating PV neurons at varying durations and frequencies, as described in the preceding section. Two-photon images of an intra-cortical sections illustrate the successful expressions of optogenetic and calcium reporter proteins at both approximately 200 μm (layer 2) and 400 μm (deeper layer 3) below the cortical surface (Figure 5A-B). Arterial diameter measurements from layer 2, depicted in Figure 5C, revealed immediate vasodilation following 1-second optogenetic stimulation (5-Hz trains of 10-ms light pulse) of PV neurons. In contrast, measurements from deeper layer 3, illustrated in Figure 5D, indicated delayed vasodilation following 4-second optogenetic stimulation (5-Hz trains of 10-ms light pulse) of PV neurons. We plotted the peak time of the maximum vascular responses following stimulation as a function of cortical depth (Figure 5E). The trend showed an increase in peak time with increasing depth. A linear regression model derived a line with a slope of -0.026 μm/s, an intercept of 10.145 s, and a correlation coefficient of 0.40 between depth and peak time.

**Figure 5.**
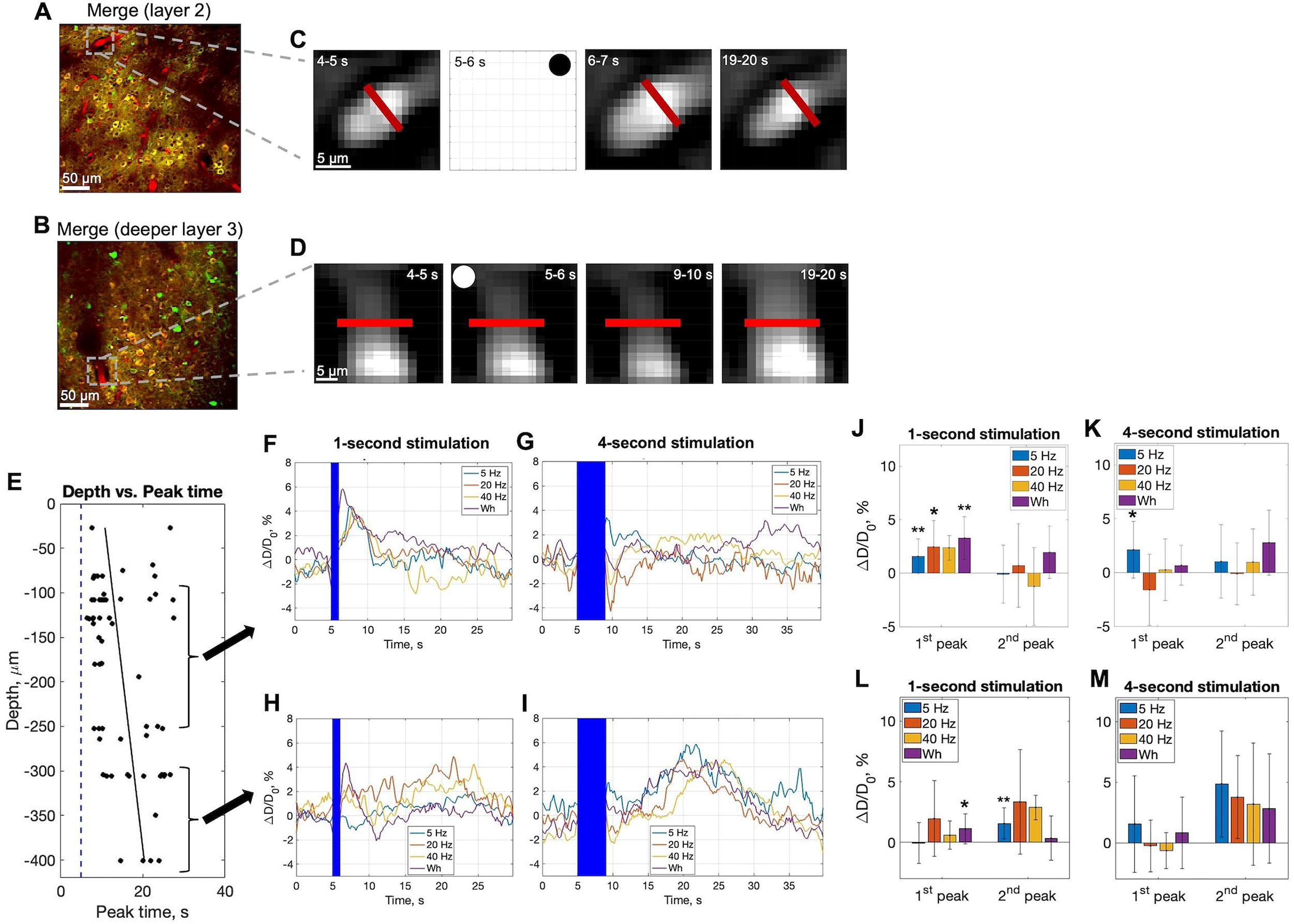
Optogenetic stimulation of PV neurons induces depth-dependent vascular response. **A.** 2P microscopy color merge image of S1 intra-cortical section at ∼250 μm and **B**. at ∼400 μm depth. **C**. Zoom-in frames of arteries located in the top-left corner of image in A and **D**. of artery located in the bottom-left corner of image in B at pre-stimulation (4-5 s), during stimulation (5-6 s), immediately after stimulation (6-7 s/9-10 s), and post-stimulation (19-20 s). Red bar indicates the region where the diameter was measured. Black and white circles indicate the frames during stimulation period. **E.** Peak time of vessel diameter change as a function of cortical depth with a fitted linear regression model (slope=-0.026 μm/s, intercept=10.145 s, correlation coefficient=0.396). Dotted line represents the beginning of stimulation (5 s). **F**. Time courses of arterial diameter changes in superficial layers in PV-cre mice expressing ChR2 in 1-s and **G.** 4-s stimulation experiments; **H**. in deeper cortical layers in 1-s and **I.** 4-s stimulation experiments. **J**. Summary of average arterial diameter changes in superficial layers at different frequencies in 1-s (n=3-13 animals) and **K**. 4-s stimulation experiments (n=3-7 animals); **L**. in deeper layers at different frequencies in 1-s (n=3-8 animals) and **M**. 4-s stimulation experiments (n=3-6 animals). Error bars denote standard deviation. Significant differences from baseline are denoted by * for p<0.05 or by ** for p<0.01.

To examine this depth-dependent effect, we grouped responses into superficial for those between 50 to 200 μm and deeper for those between 250 to 400 μm. On average, 1-second optogenetic stimulation of PV neurons in superficial layers robustly induced fast vasodilation (Figure 5F). The 1^st^ peak in vessel diameter change was similar across frequencies, with amplitudes of 1.6 ± 1.6%, 2.5 ± 2.5%, and 2.4 ± 1.1% at 3.2 s, 2.8 s, and 2.6 s after stimulus onset for 5 Hz, 20 Hz, and 40 Hz, respectively (Figure 5F, J). Statistical significance was observed for 5 Hz and 20 Hz (n=12, p<0.01; n=8, p<0.05; Figure 5J). In contrast, in 4-second optogenetic stimulation of PV neurons in superficial layers only 5-Hz stimulation induced fast vasodilation: 2.1 ± 2.6% at 5.2 s after stimulus onset (Figure 5G, K). Interestingly, after 20 Hz and 40 Hz stimulation, we observed fast vasoconstriction (Figure 5G): -1.6 ± 3.3% and 0.3 ± 3.3% at 5.9 s and 5.7 s after stimulus onset for 20 Hz and 40 Hz, respectively, but it did not reach significance (n=3, p=0.5; n=3, p=0.25; Figure 5G, K). Linear mixed-model analysis demonstrated no significant relationship between the 1^st^ peak amplitude and optogenetic stimulus frequency for 1-second (p=0.71) or 4-second stimulations (p=0.87). No significant 2^nd^ peaks were observed in stimulations of any duration. These results demonstrate that short (1 s) optogenetic stimulation of PV neurons in superficial layers (layer 1-2) induces fast arterial vasodilation; however, long optogenetic stimulation (4 s) at higher frequencies might cause vasoconstriction.

In contrast, optogenetic stimulation of PV neurons in deeper layers did not induce robust fast vasodilation (Figure 5H-I). Instead, it produced a delayed slow vasodilation, especially for 4-second stimulation (Figure 5I). In 1-second stimulation, the 2^nd^ peak in vessel diameter change was 1.6 ± 1.3%, 3.3 ± 4.3%, and 2.9 ± 1.0% at 14.9 s, 11.9 s, and 19.6 s after stimulus onset for 5 Hz, 20 Hz, and 40 Hz, respectively (Figure 5H, L). In 4-second stimulation, the 2^nd^ peak in vessel diameter change was 4.9 ± 4.4%, 3.8 ± 3.4%, and 3.2 ± 5.0% at 20.8 s, 15.1 s, and 17.5 s after stimulus onset for 5 Hz, 20 Hz, and 40 Hz, respectively (Figure 5I, M). Statistical significance was observed for the 2^nd^ peak in 5-Hz 1-second optogenetic stimulation (n=8, p<0.01; Figure 5L). Linear mixed-model analysis demonstrated no significant relationship between the 2^nd^ peak amplitude and optogenetic stimulus frequency in 4-second stimulation (p=0.87). These findings indicate that prolonged (4 s) optogenetic stimulation of PV neurons in deeper layers (layer 3) induces a delayed slow arterial vasodilation, aligning with the CBF changes measured with LDF in Figure 2B-D and suggesting that this effect persists into even deeper layers. Importantly, these results suggest that the vascular regulation by PV neurons exhibits depth-dependent characteristics.

To confirm that the observed changes in arterial diameter were solely due to optogenetic stimulation of ChR2, we conducted experiments in mice lacking ChR2 (RGECO1, PV-mCherry). We observed no changes in arterial diameter following any of the optogenetic stimulation parameters tested (Supplemental Figure 4).

Both 1-second and 4-second whisker stimulations elicited typical fast vasodilation at both superficial and deeper layers. A peak observed behind the stimulation bar in the 4-second whisker stimulation experiment (Supplemental Figure 5) aligned with the 1^st^ peak in 1-second whisker stimulation and was disregarded for consistency with optogenetic experiments. For 1-second stimulation at superficial layers, the 1^st^ peak was 3.3% ± 2.0% at 2.2 s and the 2^nd^ peak was 2.0% ± 2.4% at 13.0 s after stimulus onset (Figure 5F, J). In the case of 4-second stimulation, the 1^st^ peak was 0.7 ± 1.9% at 6.5 s, and the 2^nd^ peak was 2.8 ± 3.0% at 25.1 s after stimulus onset (Figure 5G, K). Statistical significance was observed for the 1st peak in 1-second stimulation (n=10; p<0.01; Figure 5J). For 1-second stimulation at deeper layers, the 1^st^ peak was 1.1 ± 1.2% at 2.1 s and the 2^nd^ peak was 0.4 ± 1.8% at 12.6 s after stimulus onset (Figure 5H, L). In the case of 4-second stimulation, the 1^st^ peak was 0.8 ± 3.0% at 5.4 s, and the 2^nd^ peak was 2.8 ± 4.5% at 15.3 s after stimulus onset (Figure 5I, M). Statistical significance was observed for the 1^st^ peak in 1-second stimulation (n=5; p<0.05; Figure 5L). Although the 2^nd^ peak of the whisker response was not statistically significant, average response amplitudes over superficial and deeper layers were similar to amplitude trends of the 2^nd^ optogenetic response peak, reinforcing the notion that this delayed or sustained response is likely due to PV neuron activity. Overall, these results indicate that neurovascular mechanism during whisker stimulation in superficial layers is different from that in deeper layers.

### Ongoing activity of PV+ neurons actively modulates spontaneous arterial fluctuations

Finally, we explored the contribution of ongoing activity of PV+ and PV-neurons to spontaneous arterial fluctuations. We recorded population neuronal activity across cortical depth using RCaMP1a for about 30 minutes, while simultaneously imaging penetrating arterioles as done above. Examples of RCaMP1a changes in single PV+ and PV-cells are presented in Figures 6A-B, and their corresponding arteriole diameter changes are shown in Figures 6C-D (red trace). Figure 6E demonstrates example HRFs obtained by convolving a gamma function with RCaMP1a changes in PV+ (green trace) and PV-(purple trace) cells during ongoing activity. The HRF from a PV+ cell appeared wider than that from a PV-cell. Next, we estimated vascular activity (Figure 6C-D, black trace) by convolving the RCaMP1a signals (Figure 6A-B) with corresponding HRFs (Figure 6E). In this particular example, the Pearson correlation coefficient between measured and estimated vessel diameter change signals was higher for the PV+ cell (r=0.37, Figure 6C) compared to the PV-cell (r=0.33, Figure 6D).

**Figure 6.**
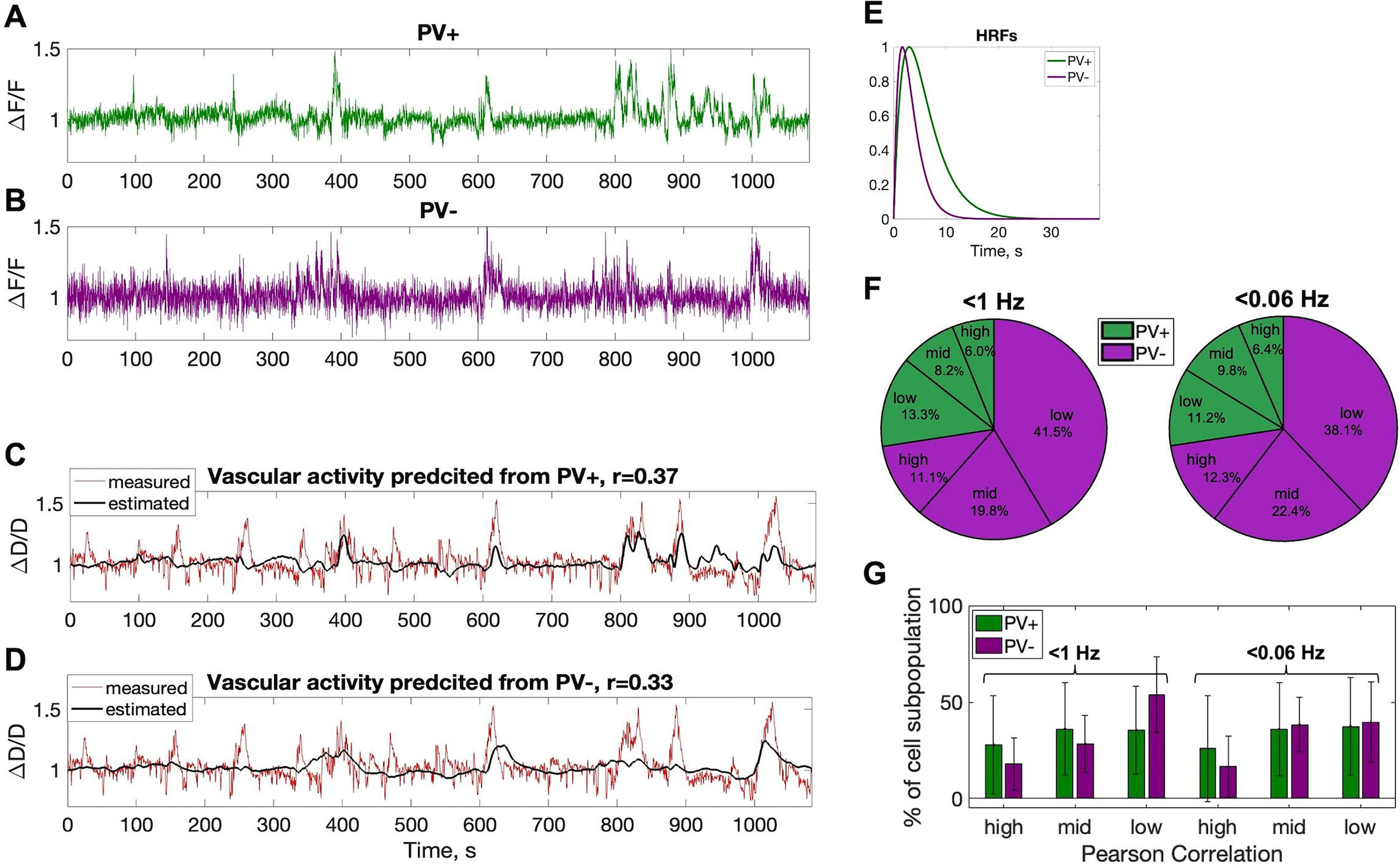
Ongoing activity of PV+ neurons actively modulates spontaneous arterial fluctuations. **A.** Time courses of RCaMP1a changes during ongoing activity in PV+ cell and **B**. PV-cell. **C**. Example time courses of measured spontaneous vessel diameter changes (red trace) and those estimated from the PV+ cell activity in A (black trace). **D.** Same measured spontaneous vessel diameter changes (red trace) and an example of those estimated from the PV-cell activity in B (black trace). **E.** HRFs obtained from convolving gamma function with RCaMP1a changes in PV+ (green trace) and PV-(purple trace) cells during ongoing activity. **F.** Summary of correlation values (high, mid, low) between measured and estimated spontaneous arterial diameter changes across cortical layers. The first group (<1 Hz) displays correlations between predicted and measured changes filtered to 1 Hz, while the second group (<0.06 Hz) shows correlations filtered to 0.06 Hz. Green color pie slices depict the percentage of PV+ cells (254 cells, ∼25%), and purple slices depict the percentage of PV-cells (751 cells, ∼75%) relative to the total number of cells (1005 cells). **G**. Green bars represent the percentage of PV+ cells within PV+ cell subpopulation, while purple bars represent the percentage of PV-cells within PV-subpopulation.

To determine which fraction of neuronal population predicts vascular activity, we categorized estimated vessel diameter change signals into three correlation groups with measured signals: higher (>0.2), mid-level (0.1-0.2), and lower (<0.1) correlation. The lower correlation group was established based on a p-value near significance threshold of 0.05 (p=0.037, 2000 data points). We also analyzed filtered slow vascular activity (0.06 Hz) to examine neuronal contributions. An example of this filtered activity and corresponding HRFs is shown in Supplemental Figure 6. From 22 recordings in 6 animals, we identified a total of 1005 cells: 254 (∼25%) were PV+ cells, and 751 cells (∼75%) were PV-cells. Note that in this analysis, PV+ cells were identified by YFP expression and responsiveness to optogenetic stimulation, while PV-cells were remaining cells visible in the field-of-view. The fractions of PV+ and PV-cells relative to the total cell number for each category are plotted in Figure 6F. For the higher (>0.2) correlation group, about 6.0% and 11.1% of cells highly predicting fast vessel diameter changes (<1 Hz) and 6.4% and 12.3% of cells highly predicting slow vessel diameter changes (<0.06 Hz) were PV+ and PV-cells, respectively. Although more PV-cells predicted vascular activity in absolute numbers, a greater fraction of PV+ cells predicted this activity at mid and high levels when considering the percentage values shown in Figure 6F relative to each cell subpopulation. At the mid-level, the PV+ cell fraction was around 33%, compared to 26% for PV-cells. At the high level, the PV+ cell fraction was about 24%, while the PV-cell fraction was around 15%. The fractions of PV+ within PV+ cell subpopulation and PV-cells within the PV-cell subpopulation for each category are plotted in Figure 6G. Approximately 28% of PV+ cells and 17.9% of PV-cells highly predicted fast vascular activity, while 25.9% of PV+ cells and 16.5% of PV-cells highly predicted slow vascular activity. Notably, more PV+ cells highly predicted both fast and slow vascular activity compared to PV-cells. These results suggest that during ongoing activity, a larger fraction of PV+ cells modulate spontaneous arterial fluctuations compared to PV-cells.

## Discussion

In this study, we explored the relationship between the activity of PV neurons and cerebrovascular dynamics during evoked and ongoing activity. First, we replicated our previous experiments [19] measuring CBF responses evoked by optogenetic stimulation of PV neurons in awake mice using LDF. Our experiments with different stimulation parameters showed that the CBF response had a stronger dependence on the frequency rather than the pulse width of optogenetic stimuli. Unlike whisker stimulation, which resulted in a rapid CBF increase, prolonged optogenetic stimulation of PV neurons (4 s vs. 1 s) resulted in a delayed, slow CBF increase as previously reported by our group and others [19–20]. This slower CBF response was also present in whisker stimulation experiments as a slow secondary peak and could be explained by the activation of PV neurons using a linear convolution model. These results suggest that PV neurons contribute (or produce) the slower sustained sensory-evoked hemodynamic response. We then used 2P Ca^2+^ imaging to examine the impact of optogenetic stimulation on PV neuron activation and network activity. Our observations revealed robust activation of PV+ neurons and inhibition of select PV-neurons during short and prolonged optogenetic stimulation. The rise of Ca^2+^ fluorescence in PV+ cells during short optogenetic stimulation was independent of stimulus frequency, whereas longer optogenetic stimulation exhibited a dependence on frequency. In those experiments we also observed the inhibition of other neurons, manifesting as a drop in Ca^2+^ fluorescence and labeled as PV-, that was greater during prolonged optogenetic stimulation. More importantly, concurrent 2P imaging of arteriolar diameter showed depth-dependent dynamics. Optogenetic activation of PV neurons produced a rapid vasodilation at low stimulation frequency and vasoconstriction at high frequencies over superficial layers (1-2), while also producing a delayed and slow vasodilation at deeper layers (3-4). Experiments conducted while recording ongoing resting-state activity showed that both PV+ and PV-populations similarly explain nearby changes in arterial diameter, but a larger fraction of PV+ cells explain arteriolar changes. Overall, our findings show complex vasodynamics with PV neuron activation that show depth-dependent characteristics dominated by slow vascular changes.

We assessed changes in local vascular tone using 2P to evaluate neuronal population contributions to hemodynamic responses across layers 1 to 4. In the deeper imaged layers, short and prolonged optogenetic stimulation of PV neurons induced delayed, slow arterial vasodilation in a stimulus dependent manner. This vasodilation resembled the CBF increase measured by LDF in similar experiments conducted in awake and anesthetized mice [19–20]. This agreement between LDF and 2P measurements indicates that slow vascular response is likely present in even deeper layers since our LDF’s measurements are sensitive to deeper locations, spanning a volume of about 1 mm^3^ (780-nm light). More importantly, we did not observe post-stimulation bounce back of neuronal activity from sustained network inhibition with the stimulation parameters we used, reinforcing the notion that this is a vascular response feature of PV neuron activity. This slower vascular response was also present at these deeper layers in prolonged whisker stimulation experiments as a slow secondary peak, also indicating that this delayed or sustained response is likely due to PV neuron activity. Additionally, preliminary experiments done using optical imaging of intrinsic signals (OIS) showed that high frequency optogenetic stimulation induced a delayed, gradual increase in CBV in the tissue and pial vessel regions (Supplemental Figure 3D, H), reflecting contributions from both superficial and deeper cortical layers. These findings strongly suggest that PV neurons induce a slow increase in CBF in deeper layers, and are consistent with previous reports that have explored examined hemodynamic responses to longer periods of optogenetic PV neuron activation [22]. Interestingly, depth-dependent fMRI measurements made in that study in ketamine-xylazine anesthetized mice showed the opposite effect with a sustained vasoconstriction across layers that persisted about 10 s with optogenetic stimulation for 20 s, that was followed by superficial then deeper vasodilation. However, optogenetic stimulation experiments conducted in awake mice did not produce the slow vasodilation despite the similarity in experimental conditions. This might be due to the relatively long stimulation period (20 s at 20 Hz) used in that study. It is unclear if this effect would disappear with the elimination of PV neuron activity. Optogenetic experiment using inhibitory opsins in PV cells could provide insight. We speculate that deactivating PV cells will eliminate the slow secondary vascular response to whisker stimulation.

In agreement with previous studies, we found that optogenetic stimulation of PV neurons induced rapid arterial vasodilation in superficial layers, similar to the first peak typically observed in whisker stimulation experiments. This was also observed in a pilot OIS experiment, which is more sensitive to hemodynamic changes in superficial layers (Supplemental Figure 3). Optogenetic stimulation at 5 Hz for 4 s increased CBV in large pial vessels and tissue (Supplemental Figure 3B). The time profile of this rapid arterial vasodilation in superficial layers matched the time profile of activated PV neural activity observed with 2P Ca^2+^ imaging. Additionally, in superficial layers, prolonged high-frequency stimulation of PV neurons led to vasoconstriction in many of our experiments although on average this response was not as prominent. This effect was also seen in our pilot OIS imaging experiment conducted at higher frequency (Supplemental Figure 3H). The time profile of this arterial vasoconstriction in superficial layers matched the time profile of suppressed excitatory neural activity observed with 2P Ca^2+^ imaging. These results are consistent with findings observed in both anesthetized mice and in awake mice using OIS imaging [21–22].

We speculate that the vascular response induced by optogenetic stimulation of PV neurons results from slow nitric oxide (NO) release, a potent vasodilator, released by neurons expressing NOS1. This mechanism was examined by Vo, et al., under light anesthesia where they observed strong slow vasodilations [22]. This mechanism is appealing because, although PV cells are present in superficial layers of somatosensory cortex, they are most densely populated in cortical layers 4 and 5, as shown in Figure 7 compiled from the Allen Mouse Brain Atlas [12, 23, 30–31]. A subset of PV neurons can release the neuropeptide substance P (SP; also known as tachykinin or Tac1; Figure 7C) [34], which binds to the SP receptor (Tacr1; Figure 7D) and is co-expressed and known to depolarize type I NOS neurons (Figure 7E) [32, 34–35]. Our group previously showed that Tacr1 neurons mediate CBF by relaxing mural cells, similar to whisker stimulation [36]. The delayed vascular response to PV neuron stimulation would then result from the release and slow dispersion of SP to Tacr1 neurons. This involvement is likely necessary since simulations of NO diffusion and degradation in tissue suggest that NO production needs to be in close proximity (on the order microns to tens of microns) for a noticeable dilatory effect [33]. Type II NOS neurons are numerous in superficial layers and a subset have been reported to also co-express PV [29–32]. Such that upon optogenetic activation, NO might be produced by activated type II NOS-PV cells near pial vessels, explaining the quicker vascular response in superficial layers.

**Figure 7.**
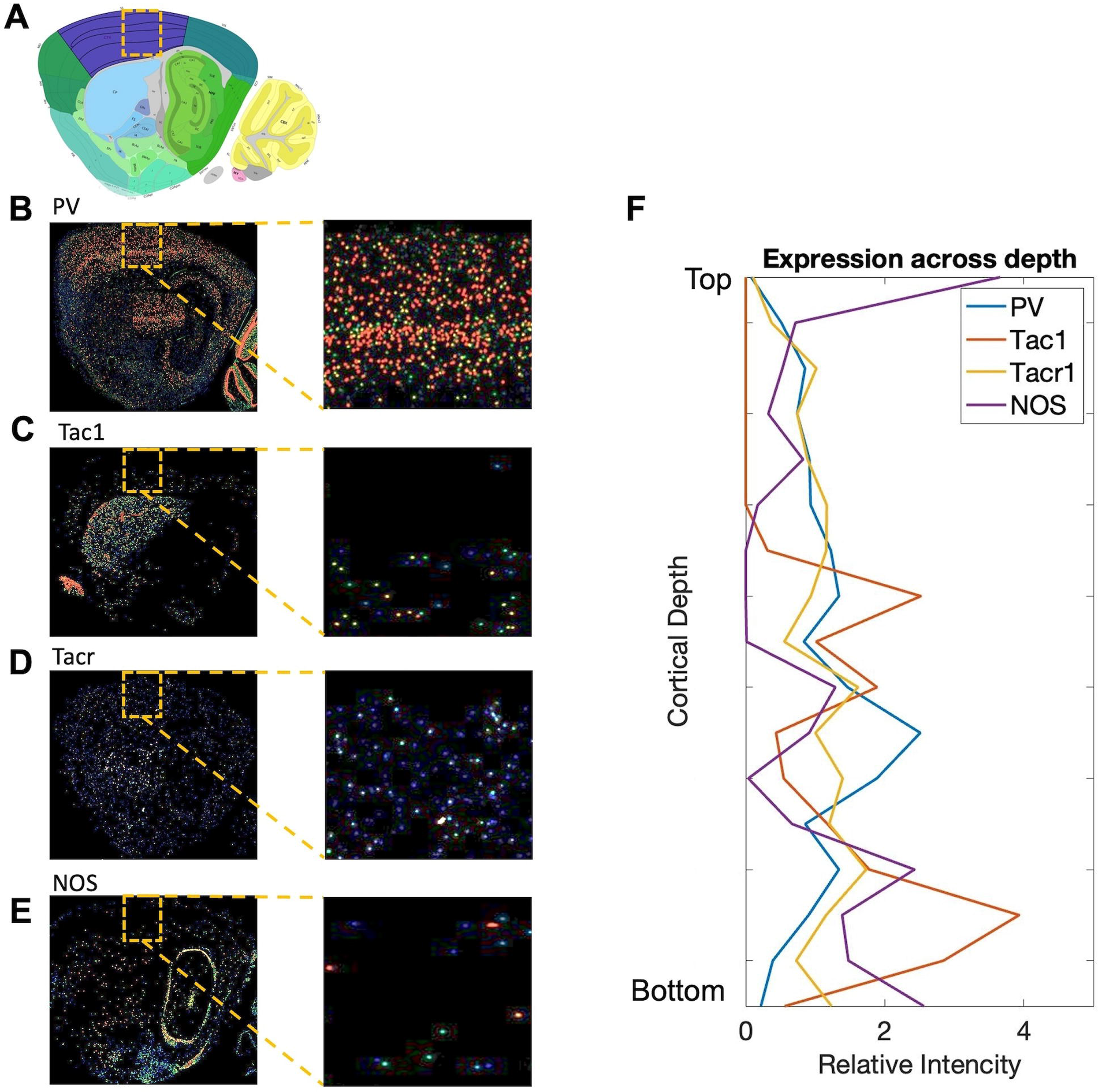
Gene expression across mouse cortical depth. **A**. Sagittal section of the mouse brain from the Allen Reference Atlas – Mouse Brain, corresponding to the slice positions in B-E [29]. The purple area indicates the somatosensory areas. **B**. Expression of PV (Allen Mouse Brain Atlas, https://mouse.brain-map.org/experiment/show/75457579), **C**. Tac1 (https://mouse.brain-map.org/experiment/show/1039), **D**. Tacr1 (https://mouse.brain-map.org/experiment/show/1296), and **E**. NOS (https://mouse.brain-map.org/gene/show/17892) in the adult mouse brain [30–31]. The second column displays zoom-in images from the orange dotted rectangle regions in the first column images. **F**. Gene expression intensity plotted as a function of cortical depth.

It is known that strong inhibitory input from PV-containing basket cells suppresses the activity of principal cells, producing a synchronized gamma rhythm [12, 23–24, 37–38]. Given that PV neurons can regulate CBF [13, 19–22], we investigated their role in modulating spontaneous arterial fluctuations. Correlation analysis between measured and estimated vascular signals revealed that while both PV+ and PV-populations actively modulate fast and slow arterial fluctuations, a higher fraction of PV+ cells influences spontaneous arterial fluctuations compared to PV-cells. Resting state is especially relevant because it lets us observe how PV neuron activity naturally influences or correlates with local vascular changes, unlike the forced and unnatural activation of these neurons. However, it is challenging to attribute correlations solely to PV neuron activity in awake mice due to the high levels of overall activity. Despite this, we anticipate that slower contributions to vascular responses would be more pronounced in deeper layers, while faster contributions would be more superficial. Our findings suggest that slow vascular fluctuations may result from PV neuron activity, supporting previous resting-state fMRI studies. We speculate that vascular tone modulation by PV+ cells is stronger at deeper layers, as evidenced by a higher correlation between PV-estimated and measured arterial diameter changes, and a greater number of PV+ cells contributing to arterial tone modulation at greater depths (Supplemental Figure 7). It would be interesting to determine whether resting-state layer fMRI experiments in humans also show preference towards deeper layers as opposed to superficial layers.

While our findings indicate that PV neuron activation induces depth-dependent vascular responses, there are several limitations to our study. We focused exclusively on the whisker somatosensory cortex and did not investigate other brain regions, which might exhibit different depth structures and vascular responses. Additionally, our ongoing activity analysis is correlational and does not establish causality. Future studies employing pharmacological interventions or chemogenetics are needed to determine the impact of PV neuron dysfunction on spontaneous vascular fluctuations. Our group is currently pursuing these approaches.

## Acknowledgements

We thank Dr. Ping Wang and Xiaoling Yang for their technical support. We also thank Dr. Karl Deisseroth and the University of North Carolina Viral Vector Core for generously providing and facilitating access to the optogenetic constructs used in this work.

## Author contributions

Conceptualization, A.R. and A.L.V.; methodology, A.R., M. F., T.D.Y.K., and A.L.V.; investigation, A.R. and A.L.V.; formal analysis, A.R. and A.L.V.; writing, A.R. and A.L.V.; supervision, A.L.V.

## Supplemental material

Supplemental material for this paper can be found at the journal website: http://journals.sagepub.com/home/jcb.

## References

1. Spencer KM, Nestor PG, Niznikiewicz MA, Salisbury DF, Shenton ME, McCarley RW. Abnormal neural synchrony in schizophrenia. Journal of Neuroscience. 2003 Aug;23(19):7407–11.

2. Celone KA, Dickerson BC, Atri A, Chua EF, Miller SL, DePeau K, Rentz DM, Selkoe DJ, Blacker D, Albert MS, Sperling RA, Fischl B. Alterations in memory networks in mild cognitive impairment and Alzheimer’s disease: An independent component analysis. Journal of Neuroscience. 2006 Oct;26(40):10222–31. doi: 10.1523/JNEUROSCI.2250-06.2006.

3. Tait L, Tamagnini F, Stothart G, Brown JT, Randall AD. EEG microstate complexity for aiding early diagnosis of Alzheimer’s disease. Scientific Reports. 2020;10:17627. doi: 10.1038/s41598-020-74790-7.

4. Cho RY, Konecky RO, Carter CS. Impairments in frontal cortical γ synchrony and cognitive control in schizophrenia. Proceedings of the National Academy of Sciences. 2006 Dec;103(52):19878–83.

5. Uhlhaas PJ, Singer W. Abnormal neural oscillations and synchrony in schizophrenia. Nature Reviews Neuroscience. 2010 Feb;11(2):100–13.

6. Lustig C, Snyder AZ, Bhakta M, O’Brien KC, McAvoy M, Raichle ME, et al. Functional deactivations: Change with age and dementia of the Alzheimer type. Proceedings of the National Academy of Sciences. 2003 Dec;100(24):14504–9.

7. Greicius MD, Srivastava G, Reiss AL, Menon V. Default-mode network activity distinguishes Alzheimer’s disease from healthy aging: Evidence from functional MRI. Proceedings of the National Academy of Sciences. 2004 Mar;101(13):4637–42.

8. Logothetis NK, Pauls J, Augath M, Trinath T, Oeltermann A. Neurophysiological investigation of the basis of the fMRI signal. Nature. 2001 Jul;412(6843):150–7. doi: 10.1038/35084005.

9. Cauli B, Hamel E. Revisiting the role of neurons in neurovascular coupling. Frontiers in Neuroenergetics. 2010 Jun;2:9. doi: 10.3389/fnene.2010.00009.

10. Lecrux C, Toussay X, Kocharyan A, Fernandes P, Neupane S, Levesque M, et al. Pyramidal neurons are ‘neurogenic hubs’ in the neurovascular coupling response to whisker stimulation. Journal of Neuroscience. 2011 Jul;31(27):9836–47. doi: 10.1523/JNEUROSCI.4943-10.2011.

11. Vazquez AL, Fukuda M, Crowley JC, Kim S-G. Neural and hemodynamic responses elicited by forelimb- and optogenetic stimulation in channelrhodopsin-2 mice: insights into the hemodynamic point spread function. Cereb. Cortex. 2014 Nov;24(11):2908–19. doi: 10.1093/cercor/bht147.

12. Rudy B, Fishell G, Lee S, Hjerling-Leffler J. Three groups of interneurons account for nearly 100% of neocortical GABAergic neurons. Dev. Neurobiol.. 2011 Jan;71(1):45–61. doi: 10.1002/dneu.20853.

13. Lee JH, Durand R, Gradinaru V, Zhang F, Goshen I, Kim D-S, et al. Global and local fMRI signals driven by neurons defined optogenetically by type and wiring. Nature. 2010 Jun;465(7299):788–92. doi: 10.1038/nature09108.

14. Perrenoud Q, Geoffroy H, Gauthier B, Rancillac A, Alfonsi F, Kessaris N, et al. Activation of cortical 5-HT(3) receptor-expressing interneurons induces NO mediated vasodilatations and NPY mediated vasoconstrictions. Front. Neural Circuits. 2012 Aug;6:50. doi: 10.3389/fncir.2012.00050.

15. Urban A, Rancillac A, Martinez L, Rossier J. Deciphering the Neuronal Circuitry Controlling Local Blood Flow in the Cerebral Cortex with Optogenetics in PV::Cre Transgenic Mice. Front. Pharmacol.. 2012 Jun;3:105. doi: 10.3389/fphar.2012.00105.

16. Anenberg E, Chan AW, Xie Y, LeDue JM, Murphy TH. Optogenetic stimulation of GABA neurons can decrease local neuronal activity while increasing cortical blood flow. J. Cereb. Blood Flow Metab.. 2015 Oct;35(10):1579–86. doi: 10.1038/jcbfm.2015.140.

17. Vazquez AL, Fukuda M, Kim S-G. Inhibitory neuron activity contributions to hemodynamic responses and metabolic load examined using an inhibitory optogenetic mouse model. Cereb. Cortex. 2018 Nov;28(11):4105–19. doi: 10.1093/cercor/bhy225.

18. Lee L, Boorman L, Glendenning E, Christmas C, Sharp P, Redgrave P, et al. Key aspects of neurovascular control mediated by specific populations of inhibitory cortical interneurons. Cereb Cortex 2020;30:2452–64. 10.1093/cercor/bhz251.

19. Krawchuk MB, Ruff CF, Yang X, Ross SE, Vazquez AL. Optogenetic assessment of VIP, PV, SOM and NOS inhibitory neuron activity and cerebral blood flow regulation in mouse somato-sensory cortex. J. Cereb. Blood Flow Metab.. 2020 Jul;40(7):1427–40. doi: 10.1177/0271678X19870105.

20. Dahlqvist MK, Thomsen KJ, Postnov DD, Lauritzen MJ. Modification of oxygen consumption and blood flow in mouse somatosensory cortex by cell-type-specific neuronal activity. J. Cereb. Blood Flow Metab.. 2020 Oct;40(10):2010–25. doi: 10.1177/0271678X19882787.

21. Lee J, Stile CL, Bice AR, Rosenthal ZP, Yan P, Snyder AZ, et al. Opposed hemodynamic responses following increased excitation and parvalbumin-based inhibition. J Cereb Blood Flow Metab 2021;41:841–56. 10.1177/0271678X20930831.

22. Vo TT, Im GH, Han K, Suh M, Drew PJ, Kim SG. Parvalbumin interneuron activity drives fast inhibition-induced vasoconstriction followed by slow substance P-mediated vasodilation. Proc. Natl. Acad. Sci. U.S.A.. 2023 Apr;120(18):e2220777120.

23. Cardin JA, Carlen M, Meletis K, Knoblich U, Zhang F, Deisseroth K, et al. Driving fast-spiking cells induces gamma rhythm and controls sensory responses. Nature. 2009 Jun;459(7247):663–7. doi: 10.1038/nature08002.

24. Sohal VS, Zhang F, Yizhar O, Deisseroth K. Parvalbumin neurons and gamma rhythms enhance cortical circuit performance. Nature. 2009 Jun;459(7247):698–702. doi: 10.1038/nature07991.

25. Niessing J, Ebisch B, Schmidt KE, Niessing M, Singer W, Galuske RAW. Hemodynamic signals correlate tightly with synchronized gamma oscillations. Science. 2005 Aug;309(5736):948–51. doi: 10.1126/science.1110948.

26. Shmuel A, Leopold DA. Neuronal correlates of spontaneous fluctuations in fMRI signals in monkey visual cortex: Implications for functional connectivity at rest. Hum. Brain Mapp.. 2008 Jul;29(7):751–61. doi: 10.1002/hbm.20580.

27. Winder AT, Echagarruga C, Zhang Q, Drew PJ. Weak correlations between hemodynamic signals and ongoing neural activity during the resting state. Nat. Neurosci.. 2017 Dec;20(12):1761–9. doi: 10.1038/s41593-017-0007-y.

28. Wu Y-T, Bennett HC, Chon U, Vanselow DJ, Zhang Q, Muñoz-Castañeda R, et al. Quantitative relationship between cerebrovascular network and neuronal cell types in mice. Cell Rep 2022;39:110978. 10.1016/j.celrep.2022.110978.

29. Allen Reference Atlas – Mouse Brain [brain atlas]. Available from atlas.brain-map.org.

30. Allen Institute for Brain Science. Allen Mouse Brain Atlas [dataset]. Available from mouse.brain-map.org. Allen Institute for Brain Science. 2004.

31. Lein ES, Hawrylycz MJ, Ao N, Ayres M, Bensinger A, Bernard A, et al. Genome-wide atlas of gene expression in the adult mouse brain. Nature. 2007 Jan;445(7124):168–76.

32. Perrenoud Q, Geoffroy H, Gauthier B, Rancillac A, Alfonsi F, Kessaris N, et al. Characterization of Type I and Type II nNOS-Expressing Interneurons in the Barrel Cortex of Mouse. Front. Neural Circuits. 2012 Apr;6:36. doi: 10.3389/fncir.2012.00036.

33. Haselden WD, Kedarasetti RT, Drew PJ. Spatial and temporal patterns of nitric oxide diffusion and degradation drive emergent cerebrovascular dynamics. PLoS Comput. Biol.. 2020 Jul;16(7):e1008069.

34. Vruwink M, Schmidt HH, Weinberg RJ, Burette A. Substance P and nitric oxide signaling in cerebral cortex: anatomical evidence for reciprocal signaling between two classes of interneurons. J. Comp. Neurol.. 2001 Sep;441(4):288–301.

35. Dittrich L, Heiss JE, Warrier DR, Perez XA, Quik M, Kilduff TS. Cortical nNOS neurons co-express the NK1 receptor and are depolarized by Substance P in multiple mammalian species. Front Neural Circuits 2012;6:31. 10.3389/fncir.2012.00031.

36. Ruff CF, Anaya FJ, Dienel SJ, Rakymzhan A, Altamirano-Espinoza A, Couey J, Fukuda M, Watson AM, Su A, Fish KN, Rubio ME. Long-range inhibitory neurons mediate cortical neurovascular coupling. Cell Reports. 2024 Apr 23;43(4).

37. Buzsáki G, Draguhn A. Neuronal oscillations in cortical networks. Science. 2004 Jun;304(5679):1926–9. doi: 10.1126/science.1099745.

38. Fries P. Neuronal gamma-band synchronization as a fundamental process in cortical computation. Annu. Rev. Neurosci.. 2009;32:209–24. doi: 10.1146/annurev.neuro.051508.135603.

