## Supplemental Figure for "“Parvalbumin interneuron activity induces slow cerebrovascular fluctuations in awake mice”"

### Supplemental Figure 1

Control experiments: average CBF response from non-expressing regions in PV-cre mice

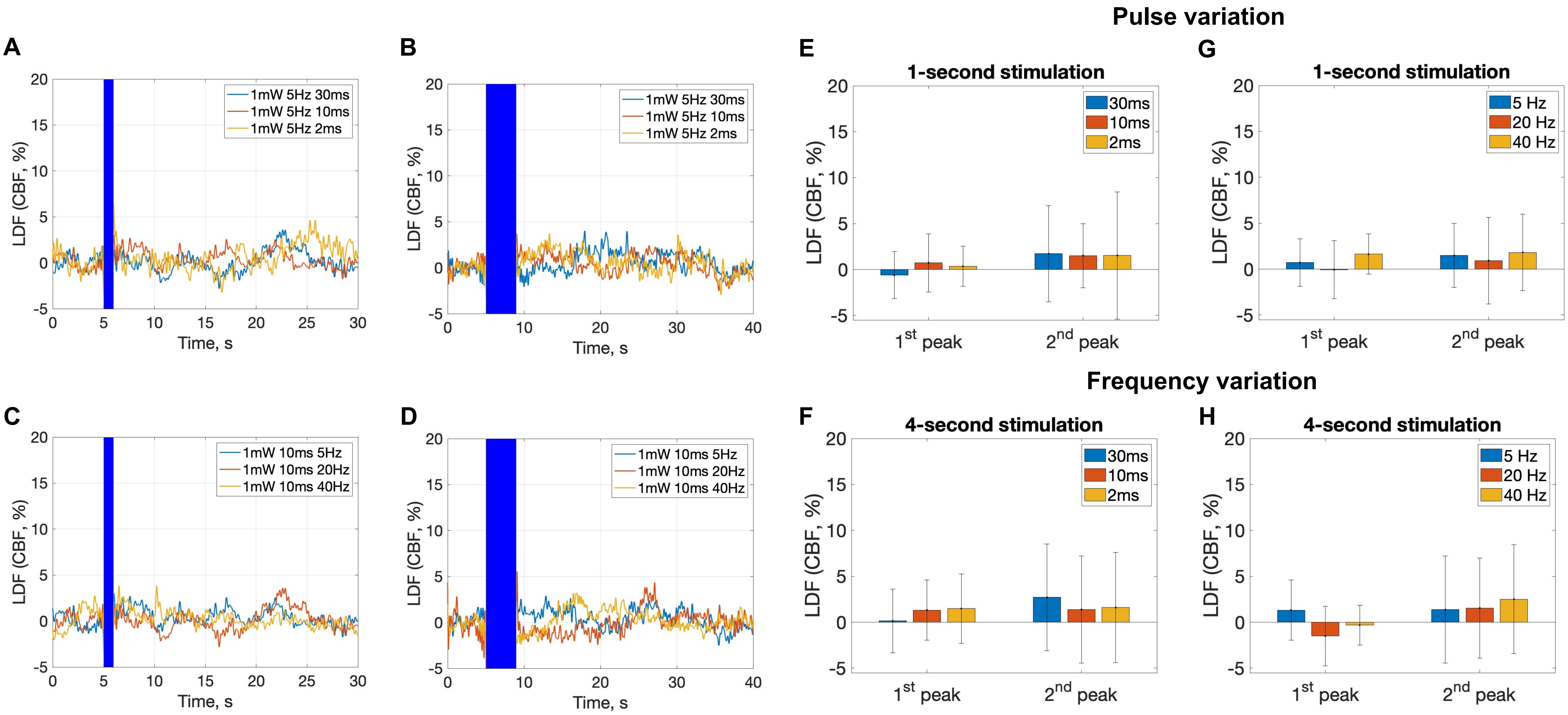

#### Supplemental Figure 2

CBF response to whisker stimulation

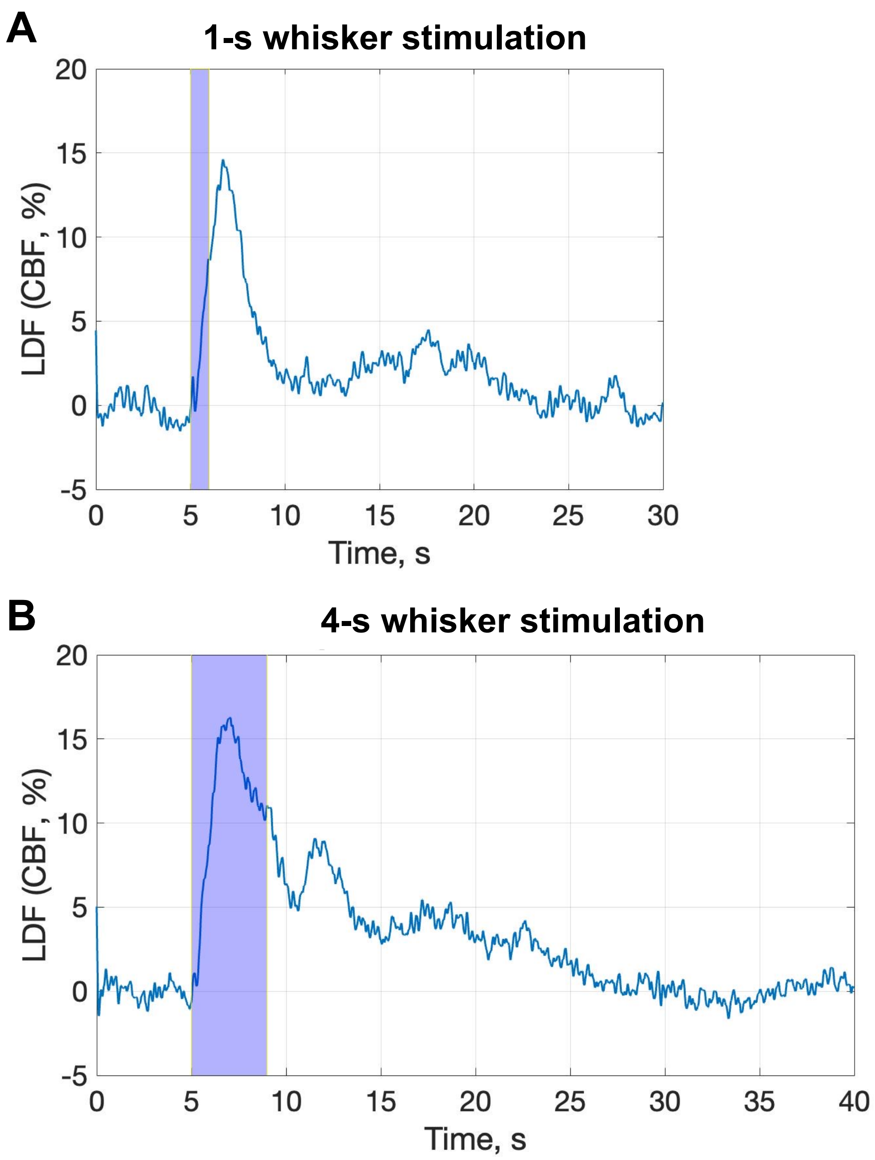

### Supplemental Figure 3

IOS imaging in PV-cre mouse expressing ChR2

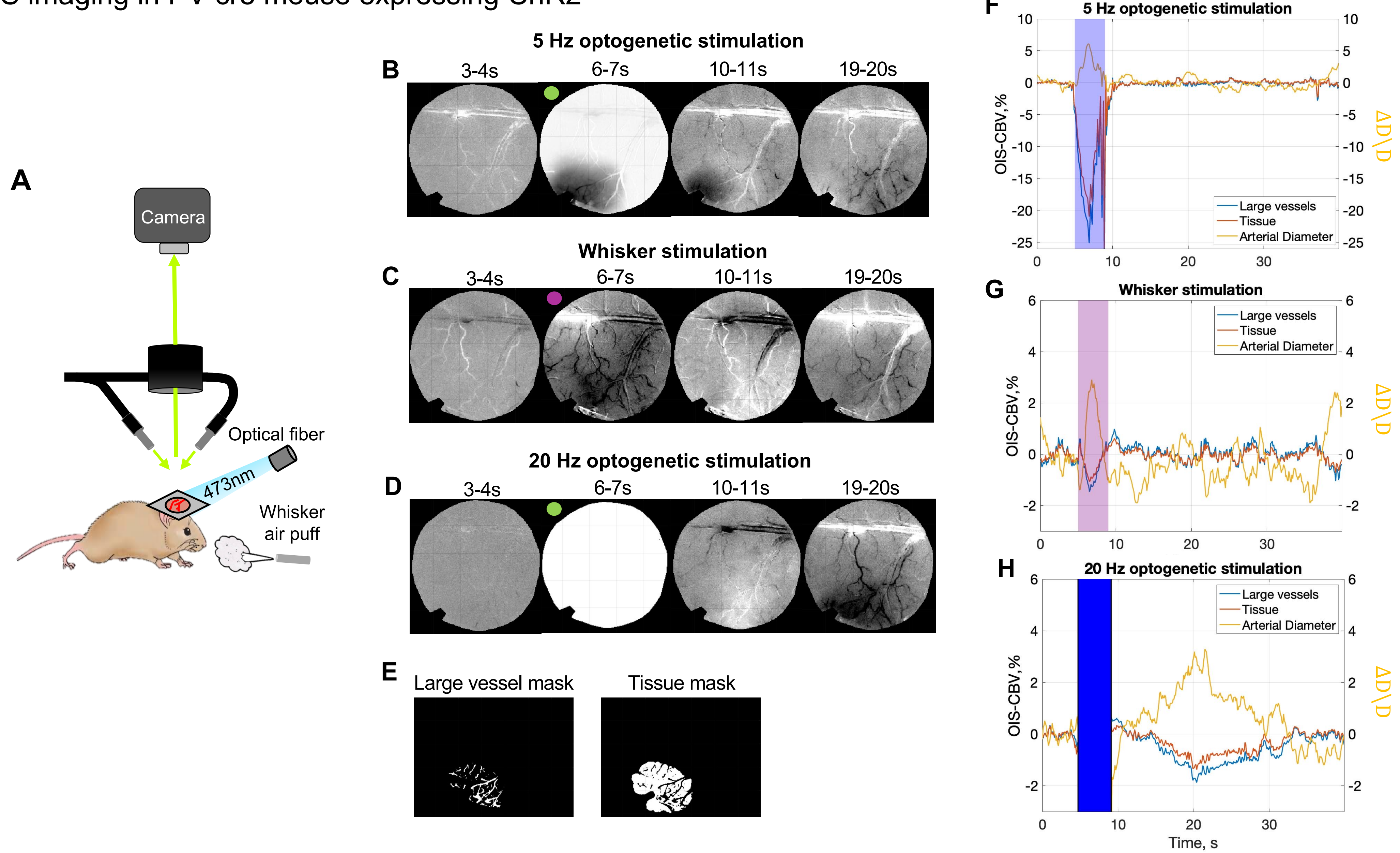

#### Supplemental Figure 4

Control experiments: average vessel diameter response in PV-cre and Thy1-RGECO1a mice lacking ChR2

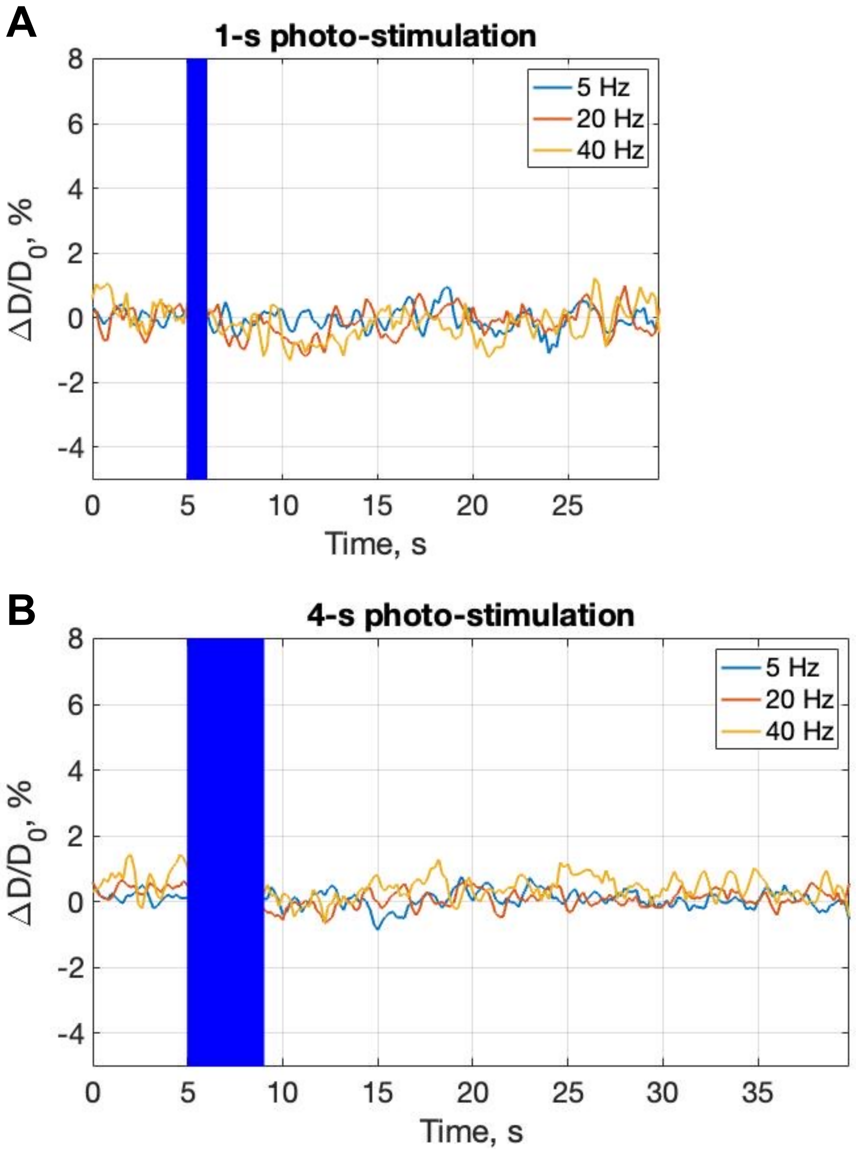

#### Supplemental Figure 5

Vessel diameter response to whisker stimulation in the surface and deeper cortical layers

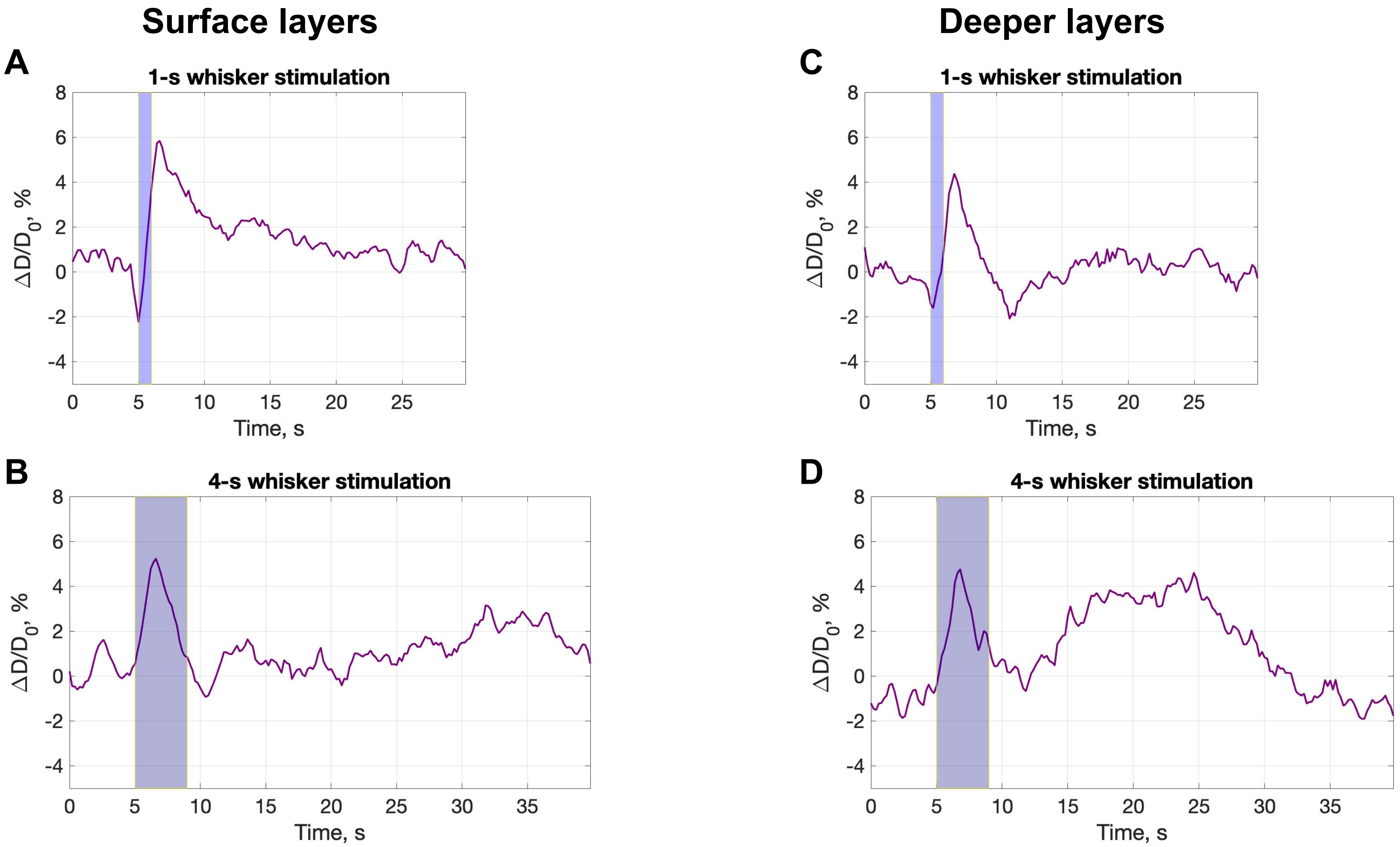

### Supplemental Figure 6

Spontaneous slow (<0.06 Hz) vascular activity and corresponding HRFs

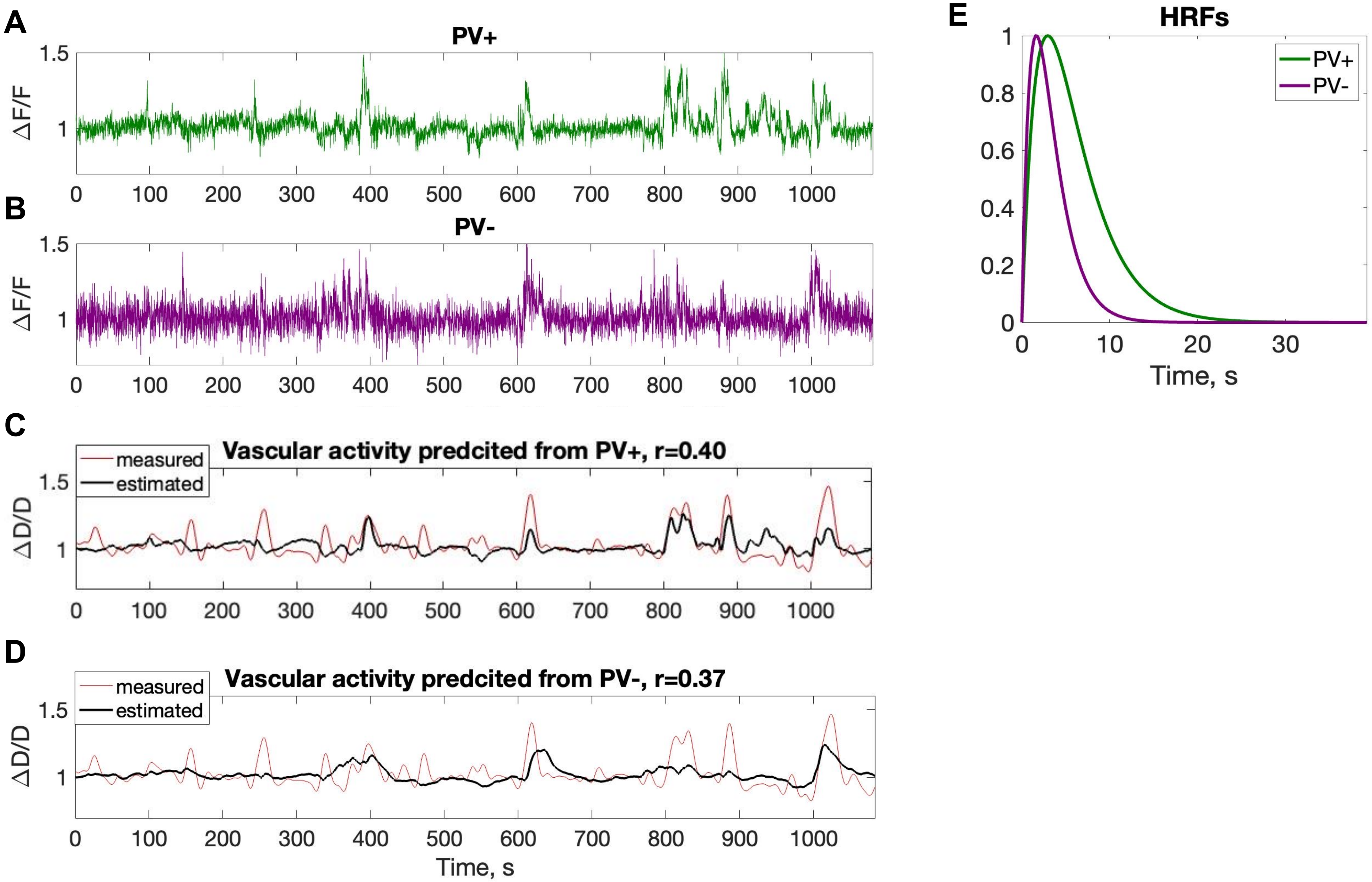

**Supplemental Figure 7**

Spontaneous neuronal and vascular activity in superficial and deeper layers

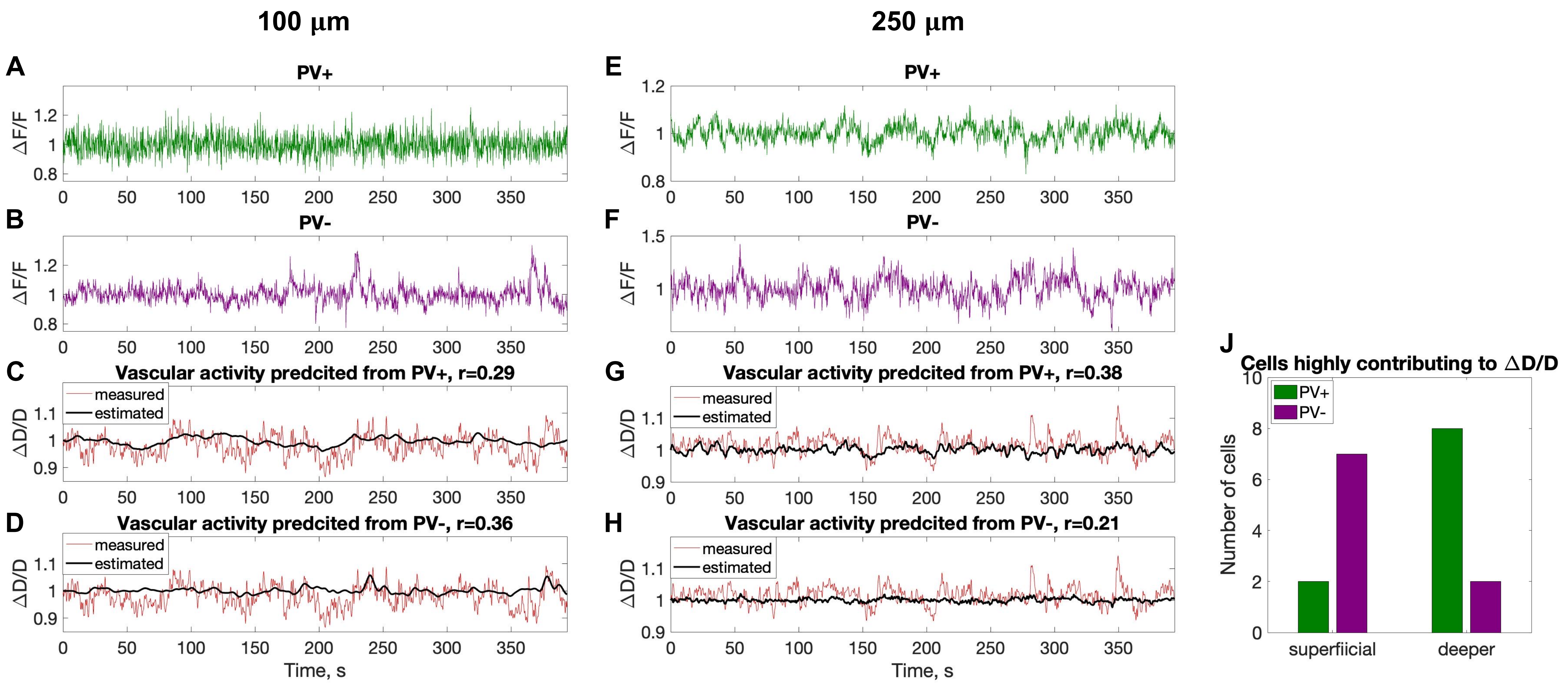
